# Transcriptomics and methylomics study on the effect of iodine-containing drug FS-1 on *Escherichia coli* ATCC BAA-196

**DOI:** 10.1101/2020.05.15.097816

**Authors:** Ilya S. Korotetskiy, Ardak B. Jumagaziyeva, Sergey V. Shilov, Tatyana V. Kuznetsova, Auyes N. Myrzabayeva, Zhanar A. Iskakbayeva, Aleksandr I. Ilin, Monique Joubert, Setshaba Taukobong, Oleg N. Reva

## Abstract

**Background:** Recent studies showed promising results on application of iodine-containing nanomicelles, FS-1, against antibiotic resistant pathogens. The effect was studied on *Escherichia coli* ATCC BAA-196.

**Materials & methods:** RNA sequencing for transcriptomics and the complete genome sequencing by SMRT PacBio RS II technology followed by genome assembly and methylomics study were performed.

**Results & conclusions:** FS-1 treated *E. coli* showed an increased susceptibility to antibiotics ampicillin and gentamicin. The analysis of differential gene regulation showed that possible targets of iodine-containing particles are cell membrane fatty acids and proteins, particularly cytochromes, that leads to oxidative, osmotic and acidic stresses. Cultivation with FS-1 caused gene expression alterations towards anaerobic respiration, increased anabolism and inhibition of many nutrient uptake systems. Identification of methylated nucleotides showed an altered pattern in the FS-1 treated culture. Possible role of transcriptional and epigenetic modifications in the observed increase in susceptibility to gentamicin and ampicillin were discussed.

**Lay abstract:** New approaches of combatting drug resistance infections are in demand as the development of new antibiotics is in a deep crisis. This study was set out to investigate molecular mechanisms of action of new iodine-containing nano-micelle drug FS-1, which potentially may improve the antibiotic therapy of drug resistant infections. Iodine is one of the oldest antimicrobials and until now there were no reports on development of resistance to iodine. Recent studies showed promising results on application of iodine-containing nano-micelles against antibiotic resistant pathogens as a supplement to antibiotic therapy. The mechanisms of action, however, remain unclear. The collection strain *Escherichia coli* ATCC BAA-196 showing an extended spectrum of resistance to beta-lactam and aminoglycoside antibiotics was used in this study as a model organism. Antibiotic resistance patterns, whole genomes and total RNA sequences of the FS-1 treated (FS) and negative control (NC) variants of *E. coli* BAA-196 were obtained and analyzed. FS culture showed an increased susceptibility to antibiotics associated with profound gene expression alterations switching the bacterial metabolism to anaerobic respiration, increased anabolism, osmotic stress response and inhibition of many nutrient uptake systems. Nucleotide methylation pattern were identified in FS and NC cultures. While the numbers of methylated sites in both genomes remained similar, some peculiar alterations were observed in their distribution along chromosomal and plasmid sequences.

---

Controlling antibiotic resistance development in bacteria is an intriguing area of research. It is generally recognized that pathogens may gain resistance to antibiotics either due to mutations modifying the target proteins or by acquisition of antibiotic resistance genes. The latter includes efflux pumps, alternative variants of target proteins, or antibiotic modifying enzymes [1]. There are various approaches to combat antibiotic resistance including new antibiotic design; drug repurposing to target alternative key molecular processes [2]; combinatorial drug therapy when several antibiotics with alternative mechanisms of action are used [3]; application of inhibitors of efflux pumps, antibiotic depleting enzymes and other proteins involved in bacterial persistence and adaptation [4]; and drug-induced reversion of selective pressures towards antibiotic susceptibility in bacterial populations by imposing a direct cost on the resistance [5]. Induction of antibiotic resistance reversion by an iodine-containing nano-molecular complex, FS-1, was demonstrated *in vitro* on multidrug resistant *Mycobacterium tuberculosis* [6-8] and *Staphylococcus aureus* [9]. FS-1 is a complex of dextrin-polypeptide ligands with iodine stabilized by coordination links between the matrix of organic molecules with triiodide, negative iodine and chloride ions, and positive lithium and magnesium ions. The general composition formula of FS-1 is [{(L_*n*_ (MeJ_3_)^+^)_*y*_ {Me(L_*m*_)J]^+^_*x*_} (Cl^−^)_*y+x+k*_], where L is the dextrin-polypeptide ligand; Me – Li/Mg ions; Cl^−^ – chloride ion; J – iodine ion; J_3_ – triiodide; *n, m, x, y* and *k* – variable integers ≥ 1. FS-1 was primarily introduced as a new therapeutic agent against multidrug resistant tuberculosis (MDR-TB) [10, 11] and currently is under the 3^rd^ round of clinical trials in Kazakhstan (www.clinicaltrials.gov, acc. NCT02607449) as a supplementary drug against MDR-TB. Further laboratory experiments demonstrated that cultivation of several multidrug-resistant bacteria *Staphylococcus aureus* [9, 12] and *Escherichia coli* [13] with this drug increased the susceptibility of these microorganisms to antibiotics. In many aspects the antibiotic resistant strains *S. aureus* and *E. coli* were found to be more convenient laboratory models to study molecular and genetic mechanisms of the therapeutic action of the iodine-containing nano-micelles rather than drug-resistant clinical isolates of *M. tuberculosis*. A research scheme of experiments on the influence of FS-1 on model antibiotic resistant microorganisms was developed and first exploited on *S. aureus* ATCC BAA-39. This research follows the same experimental scheme described in the previous publication [9].

This work presents results of whole genome and RNA sequencing of the FS-1 treated (FS) and the negative control (NC) cultures of the model multidrug resistant microorganism *Escherichia coli* ATCC BAA-196. The study was aimed at identification of genetic, transcriptional and epigenetic modifications caused by cultivation of the model microorganism with a sub-bactericidal concentration of FS-1. Possible involvement of epigenetic modifications in antibiotic resistance reversion was hypothesized during the previous studies on the multidrug resistant *M. tuberculosis* as the whole genome sequencing of the FS-1 treated culture showing an increased susceptibility to antibiotics did not reveal any significant mutations to explain this phenomenon [6]. Further studies on *S. aureus* demonstrated significant changes in patterns of epigenetically modified nucleotides in the FS-1 treated culture [9].

The strain *E. coli* ATCC BAA-196 was laboratory designed by transferring the antibiotic resistance plasmid pMG223 obtained from *Klebsiella pneumoniae* ATCC 14714 to *E. coli* J53-2 [14]. Pathogenic *E. coli* are common agents of nosocomial infections [15-17]. Outbreaks of foodborne infections associated with antibiotic resistant *E. coli* are being reported around the world with a growing frequency [18-20]. It is alarming that the drug resistant isolates of *E. coli* are also frequent in environmental habitats including water resources [21] and agricultural products [22]. Recent discoveries showed that epigenetic modifications may play an important role in *E. coli* virulence development. An example of such regulation is controlling of the expression from a pyelonephritis-associated pili (pap) operon in uropathogenic *E. coli* by methylation of a GATC site in the promoter region reported by Hernday *et al.* [23]. The operon is transcribed when the methylation occurs proximal to the promoter, and vice versa. Another role of GATC partially methylated sites is switching between alternative promoters of an operon as it was shown on an example with the *sci1* gene cluster in enteroaggregative *E. coli* [24].

Complete genome sequencing of the collection strain *E. coli* ATCC BAA-196 (CP042865) originated from *E. coli* J53-2 (K-12 isolate) confirmed clustering of this microorganism with other genomes of the *E. coli* K-12 group: NC_000913, W3110 (NC_007779) and DH10B (NC_010473). *E. coli* K-12 group represents the normal avirulent human gut microflora [25-27]. The major difference between the multidrug resistant strain BAA-196 and the avirulent *E. coli* K-12 isolates is the presence of the large plasmid (CP042866) of the *K. pneumoniae* origin comprising multiple drug resistance determinants [14]. This plasmid shows a significant sequence similarity to another virulence plasmid pKP64477b from *Klebsiella pneumonia* [28] and some other similar plasmids frequent in different species of Enterobacteriaceae. This makes *E. coli* ATCC BAA-196 an ideal model organism to study antibiotic resistance acquisition by plasmid exchange and to model the mechanisms of antibiotic resistance reversion, as the K-12 strains are among the best-studied bacteria in terms of biology, metabolism and gene regulation [29, 30]. In this paper, possible mechanisms of increasing the susceptibility of *E. coli* ATCC BAA-196 treated with FS-1 to gentamicin and ampicillin observed in experiments were investigated by comparison of gene expression profiles and patterns of methylation of chromosomal and plasmid DNA in FS and NC culture variants.

## Materials & methods

### Bacterial cultures

The model multidrug resistant microorganism *Escherichia coli* ATCC BAA-196™ producing extended spectrum beta-lactamases was obtained from the ATCC collection (https://www.lgcstandards-atcc.org/en.aspx) and kept in freezer at minus 80°C. Lyophilized culture obtained from ATCC was reactivated in two passages in Mueller-Hinton (MH) liquid medium (Himedia, India) supplemented with ceftazidime 10 μg/ml as recommended by the ATCC product sheet.

### Antibiotic susceptibility tests

Susceptibility of bacterial cultures to ampicillin and gentamicin was evaluated by serial dilution in 100 ml liquid MH medium with 0.05% resazurin (Sigma-Aldrich) using 96 well plates. Plates were inoculated with 20 μl overnight bacterial cultures diluted to 1.5×10^6^ CFU/ml and then incubated at 37°C for 24 h. Optical density (OD) was measured at 540/620 nm using Multiscan Ascent and compared to the negative control well with resazurin. OD values were transformed to CFU/ml values using resazurin calibration curve determined in advance using the conventional plate counting on Petri dishes. Each concentration level was measured in triplicate to calculate mean and standard deviation values [31].

Each antibiotic concentration shown in Fig. 1 was repeated six times to calculate the average OD values and the standard deviations.

**Fig. 1.**
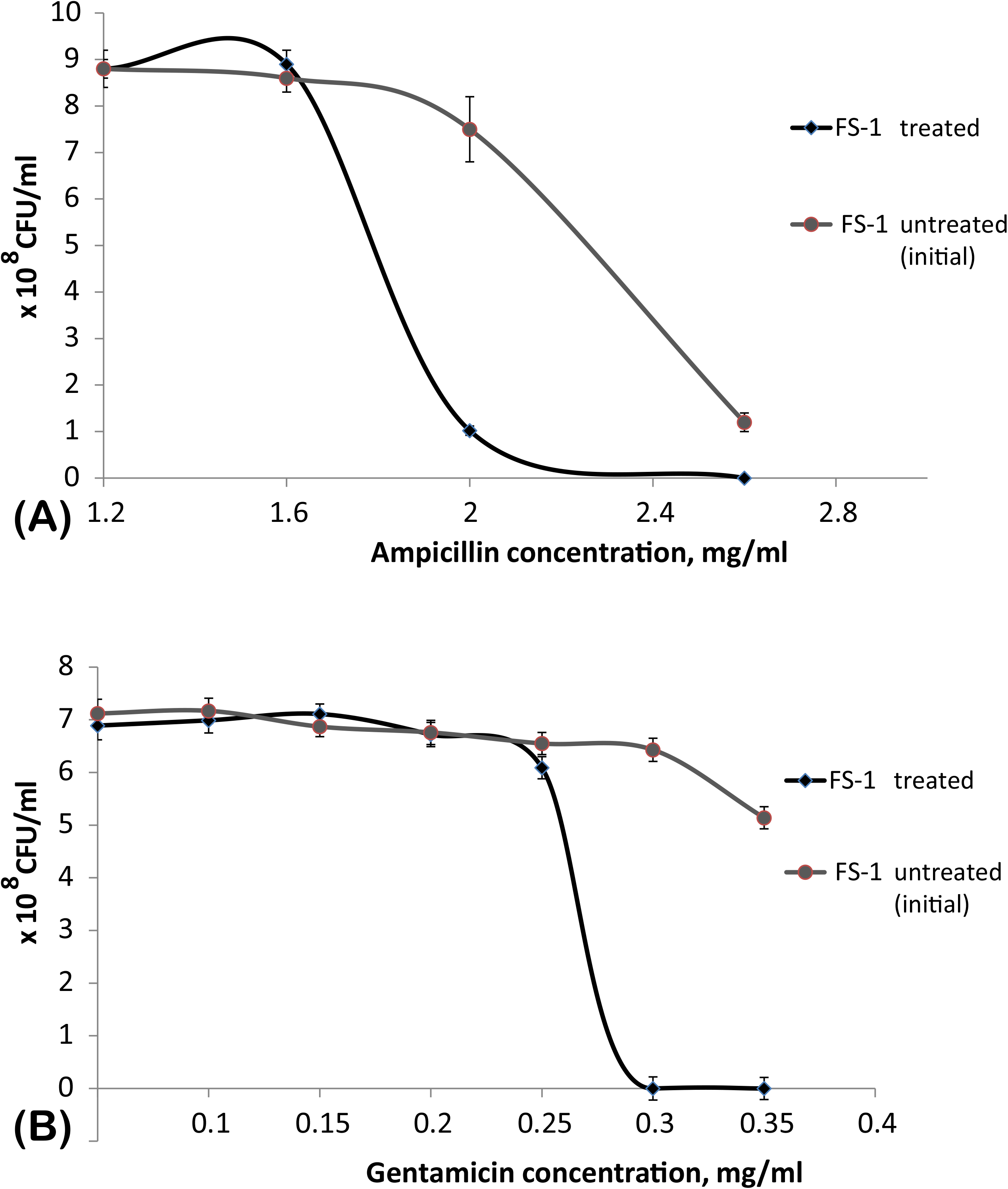
Susceptibility of the initial and FS-1 treated *E. coli* cultures to gentamicin and ampicillin. X axis depicts concentrations of antibiotics (A) ampicillin; and (B) gentamicin. Y axis shows CFU/ml values calculated based on OD values measured at 540/620 nm. OD values for every antibiotic concentration were measured in 6 wells, then the average values and standard deviations were calculated.

### Cultivation with FS-1

The experiment of cultivation of antibiotic resistant model microorganisms with FS-1 and the graphical scheme of the experiment were published before [9]. Bacteria were inoculated into test-tubes with 10 ml of liquid MH medium supplemented with FS-1 (500 μg/ml as the final concentration) that corresponds to ½ minimal bactericidal concentration (MBC) of the drug calculated previously for *E. coli*. Neutral pH of the medium was controlled. As a negative control, the culture was cultivated in the same medium without FS-1. Test-tubes were incubated at 37°C for 24 h and then 0.1 ml aliquots of the cultures were transferred to fresh tubes with the corresponding media. After 10 passages of daily re-inoculations, the experimental and control bacteria were cross-inoculated in three repetitions into tubes with FS-1 containing and FS-1 free media for further overnight incubation, followed by DNA extraction.

### HPLC detection of intracellular antibiotics

*E. coli* culture was inoculated into test-tubes with 10 ml of liquid MH medium and cultivated for 18 – 24 h at 37°C with 100 rpm shaking till achieving 5.0 OD McFarland standard units corresponding to 1.5×10^9^ CFU/ml. Cells were pelleted by centrifugation for 5 min at 5,000 rpm, supernatant was removed and the cells were washed three times and resuspended in 0.8 ml of 1,000 μg/ml stock solutions of antibiotics (amoxicillin, ampicillin and piperacillin) mixed with 0.8 ml FS-1 stock solution (1,000 μg/ml) or 0.8 ml sterile water as control samples. The final concentrations of FS-1 and antibiotics were 500 μg/ml. Each experiment with different antibiotics and controls were repeated three times to allow statistical validation of the results.

The experimental and control suspensions were incubated for 30 min at 37°C with 100 rpm shaking. Then the cells were washed three times with sterile saline and pelleted by centrifugation for 5 min at 5,000 rpm.

Cell pellets were resuspended in a 1.6 ml lysis buffer (10 mM Tris-HCl, 1 mM EDTA, 1% SDS and 6% Triton X100) and incubated for 60 min at 37°C. After incubation, tubes were frozen at −80°C for 20 min, melted at room temperature and ultrasonicated at 20 kHz ± 500 Hz for 2.5 min by SONOPULS Ultrasonic Homogenizer HD 20701 (BANDELIN, Berlin, Germany). To remove cell debris, solutions were centrifuged for 10 min at 14,000 g. Supernatant 50 μl aliquots were collected for further HPLC analysis on the Agilent 1200 HPLC system (Agilent Technologies, Santa Clara, California, USA). Samples were fractionated on a HPLC column at a flow rate 1 ml/min at room temperature (25°C) in 15 min using KH_2_PO_4_ / acetonitrile (ACN) buffer as the mobile phase. The following KH_2_PO_4_ / ACN buffers were used: 23/87 w/w (pH 5.0) for piperacillin; 5/95 w/w (pH 5.0) for amoxicillin; and 15/85 w/w (pH 2.0) for ampicillin. The characteristic peaks of eluted compounds were recorded by measuring OD values at 230 nm wavelength and compared to the values of eluted antibiotics used as standards (Supplementary Figures S1-3).

### Radiological study of FS-1 complexing with chromosomal DNA

To carry out the radiological study, isotope ^131^I was used for synthesis of FS-1. The documented radioactivity of the isotope 20 MBq/ml was controlled at the beginning of the synthesis. Radioactivity was measured by β-spectrometer Hidex 300 SL (Finland).

Bacteria were incubated in a thermo-shaker overnight on MH medium at 37°C. The culture growth was centrifuged at 5,000 g and the cells were resuspended in saline to achieve 1×10^9^ CFU/ml. Aliquots of 0.4 ml of the marked FS-1 solution (total radioactivity of an aliquot was 300 Kbq) were added to 7.6 ml of the bacterial suspension and rigorously shaken. Samples were incubated for 1 h at 37°C in a thermo-shaker, Thermomixer Comfort (Germany). After incubation, bacterial cells were washed from FS-1 by two rounds of centrifugation at 5,000 g and re-suspended in sterile saline. DNA samples were extracted from bacterial cells using PureLink Genomic DNA Kits (Publication Number: MAN0000601, Revision 2.0) following the manufacturer’s recommendations. DNA purity and concentration were controlled by NanoDrop 2000C (USA) at wavelengths 260 and 280 nm. Then the DNA samples were resolved in scintillation solute (ULTIMA GOLD LLT, PerkinElmer) to a final volume of 5 ml. The residual radiation was measured by β-spectrometer Hidex 300 SL (Finland). Experiments were repeated three times for statistical validation.

### DNA extraction and PacBio sequencing

DNA samples were extracted in parallel from three negative control and three FS-1 treated bacterial cultures using PureLink Genomic DNA Kits (MAN0000601, Revision 2. 0) following the manufacturer’s recommendations. Each DNA sample was sequenced independently.

Samples were prepared according to the guide of preparation of SMRTbell templates for sequencing on the PacBio RS II System. The samples were sequenced at Macrogene (South Korea) on SMRT Cell 8Pac V3 cells using the DNA Polymerase Binding Kit P6 and DNA Sequencing Reagent 4.0 v2 following the SMRTbell10-kb library preparation protocol. PacBio reads generated in three repeats from the NC and FS cultures were deposited at NCBI SRA database under accession numbers (SRX6842003, SRX6842011, SRX6842012) and (SRX6842007–SRX6842009), respectively.

### Genome assembly and annotation

The complete genome assembly was performed using SMRT Link 6.0.0 pipeline as published previously [13]. Two genome variants, NC and FS, were obtained by assembly of the joint pools of DNA reads of three sets of DNA samples generated from NC and FS cultures of *E. coli* BAA-196. The NC (CP042865) and FS (CP042867) chromosomes were 4,682,561 and 4,682,572 bp, respectively. The genomes also comprise large plasmids of 266,396 and 279,992 bp, for NC (CP042866) and FS (CP042868) variants, respectively. Moreover, FS-1 treated strain contains two smaller plasmids CP042869 and CP042870 of 44,240 bp and 11,153 bp, respectively. The completeness of the final assemblies was evaluated using the benchmarking universal singlecopy orthologous (BUSCO) software [32]. Genome annotation was performed using the RAST Server (http://rast.nmpdr.org/; [33]) and then manually corrected. Involvement of the predicted genes in metabolic pathways was predicted by the Pathway Tools 23.0 [34] and the EcoCyc Database [29]. Genes involved in antibiotic resistance were predicted by RGI 5.1.0 / CARD 3.0.5 Web-portal [35]. Locations of horizontally transferred genomic islands were identified and the general DNA sequence parameters such as GC-content and GC-skew were calculated by the program SeqWord Genome Island Sniffer [36] in 8 kbp sliding windows stepping 2 kbp along the genome sequence. The same program was used for identification of replication origins and termini on the bacterial chromosomes by GC-skew between the leading and lagging strands [37]. The obtained genomes were deposited in NCBI with the accession numbers CP042865-CP042866 for the NC variant of the chromosome and large plasmid of this strain; and CP042867-CP042870 for the chromosome and three plasmids of the FS variant. Raw PacBio sequences are available from the NCBI BioProject Website under the accession number PRJNA557356.

### RNA extraction and sequencing

Isolation of the total RNA from the NC and FS cultures after overnight cultivation (the mid logarithmic growth phase controlled by OD) on the respective media was performed as it was published before [9]. Shortly, the RiboPure Bacteria Kit (Ambion, Lithuania) was used according to the developer’s guidelines. The quality and quantity of the resulting RNA were determined using the Qubit 2.0 Fluorometer (Thermo Scientific, USA) and Qubit RNA Assay Kit (Invitrogen, USA). The MICROBExpress Bacterial mRNA Purification Kit (Ambion, Lithuania) was used for ribosomal 16S and 23S RNA removal from the samples. RNA libraries prepared with the Ion Total RNA Seq Kit V2 (Life Technologies, USA) were sequenced on the Ion Torrent PGM sequencer (Life Technologies, USA) with the Ion 318 Chip Kit V2. Three repetitions of RNA reads generated from the NC and FS variants were deposited at NCBI SRA database under accession numbers SRX6842006 and SRX6842002, respectively.

### Differential gene expression analysis

The differential expression was done using the R-3.4.4 software. Firstly, a reference index was built for each reference genome using the *buildindex* function available in the *Rsubreads* package (Bioconducter). For each bacterium, the obtained RNA fragments were aligned to the relevant reference genomes with the use of the “align” function. The aligned BAM files and relevant GFF annotation files were then used as input for the *featureCounts* function to obtain gene counts. The R packages DESeq2 (Bioconducter) and *GenomicFeatures* were then used in R studio for the differential expression analyses. The visualization of the volcano plots of differential expression in the studied genomes was performed by an in-house Python script using the output excel files generated by R package DESeq2. Networks of regulated genes were constructed using the Web based tool PheNetic [38] based on the regulation network available from the PheNetic Web site, which was designed for the strain *E. coli* K12 (NC_000913. 2). Pairs of homologous genes in the genomes K12 and BAA196 (Supplementary Table S1) were identified using the program GET_HOMOLOGUES [39].

### Profiling of epigenetic modifications

Epigenetic modifications of nucleotides were predicted using the standard SMRT Link DNA modification prediction protocol, *ds_modification_motif_analysis*, as it was explained in the previous publication [9]. DNA reads generated from every repetition of DNA sampling (three NC and three FS samples) were processed separately. The program calculates several statistical parameters such as IPDRatio of exceeding of the base call delay over the expectation and base call quality values to estimate the base modification (BM) scores representing the likelihood of modification of a nucleotide at the given strand and location. BM scores above 21 (*p*-value ≥ 0.01) were used to select statistically reliable sites of epigenetically modified nucleotides. Only those sites which showed BM scores ≥ 21 in all three DNA sample repeats were selected for further consideration. Average BM scores were calculated.

Base modification motifs were predicted by the program *motifMaker*. Visualization of profiles of epigenetic modifications was performed by an in-house Python script.

## Results

### Antibiotic susceptibility trials

This study was performed to evaluate the effect of iodine-containing nano-micelle drug FS-1 on antibiotic susceptibility of the reference multidrug resistant strain, *E. coli* ATCC BAA-196, which is characterized by resistance to the beta-lactam antibiotic ampicillin and the aminoglycoside antibiotic gentamicin. The experimental culture (FS) was cultivated for 10 passages with the sub-bactericidal concentration of FS-1 (500 μg/ml). In parallel, a negative control (NC) culture was cultivated for 10 passages in Mueller-Hinton (MH) liquid medium without FS-1 and antibiotics.

Minimal bactericidal concentrations (MBC) of the antibiotics were determined for FS and NC cultures by serial dilution in 96-well plates [40]. Cultivation with FS-1 reduced MBC for ampicillin from 2 – 2.6 mg/ml recorded for the culture NC to 1.6 – 2 mg/ml recorded for the culture FS (Fig. 1A). For gentamicin, these numbers were 0.3 – 0.35 mg/ml and 0.2 – 0.25 mg/ml for NC and FS cultures, respectively (Fig. 1B). This difference in susceptibility to antibiotics between the initial and FS-1 treated cultures was statistically reliable as illustrated by OD records for the plates with ampicillin and gentamicin in Fig 1.

### Influence of the treatment with FS-1 on intracellular antibiotic permeability

It was found that 30 min treatment of *E. coli* cells with FS-1 and antibiotics amoxicillin, ampicillin and piperacillin increased the intracellular concentration of antibiotics for several orders of magnitude compared to the treatment of the cells solely with antibiotics (Table 1, HPLC graphs for amoxicillin, ampicillin and piperacillin are shown in Supplementary Figures S1-3). It was hypothesized that iodine molecules released from FS-1 micelles halogenate cell wall and cytoplasmic membrane proteins and/or fatty acids disrupting their barrier functions.

**Table 1.** Intracellular concentrations of antibiotics applied on *E. coli* cells for 30 min solely and with 500 μg/ml FS-1. Relative concentrations of antibiotics were determined by HPLC.

| Antibiotic | Average concentration (mg/L) and standard deviation* |  |
| --- | --- | --- |
|  | Treatment solely with antibiotics | Treatment with antibiotics and 500 µg/ml FS-1 |
| Amoxicillin | 9.24 ± 0.05 | 1895.61 ± 5.77 |
| Ampicillin | 7.99 ± 0.12 | 716.05 ± 0.07 |
| Piperacillin | 13.76 ± 0.01 | 555.01 ± 0.22 |

\*Each experiment was repeated three times.

### Complete genome assembly and annotation

Complete genome sequences of NC and FS variants of *E. coli* ATCC BAA-196 were obtained by assembly of PacBio SMRT reads. Chromosomal sequences of NC and FS variants were 4,682,561 bp and 4,682,572 bp, respectively. Both genomes comprised the large plasmids 266,396 bp and 279,992 bp of NC and FS variants. Sequences of these plasmids were similar except for a 25,313 bp prophage insertion in the plasmid FS flanked with two copies of *insH* transposases and a deletion of nine genes including *traG* and *traF* conjugation genes. This deletion potentially can affect the efficacy of conjugation of the plasmid FS. The alignment of sequences of plasmids NC and FS is shown in Fig. 2A.

**Fig. 2.**
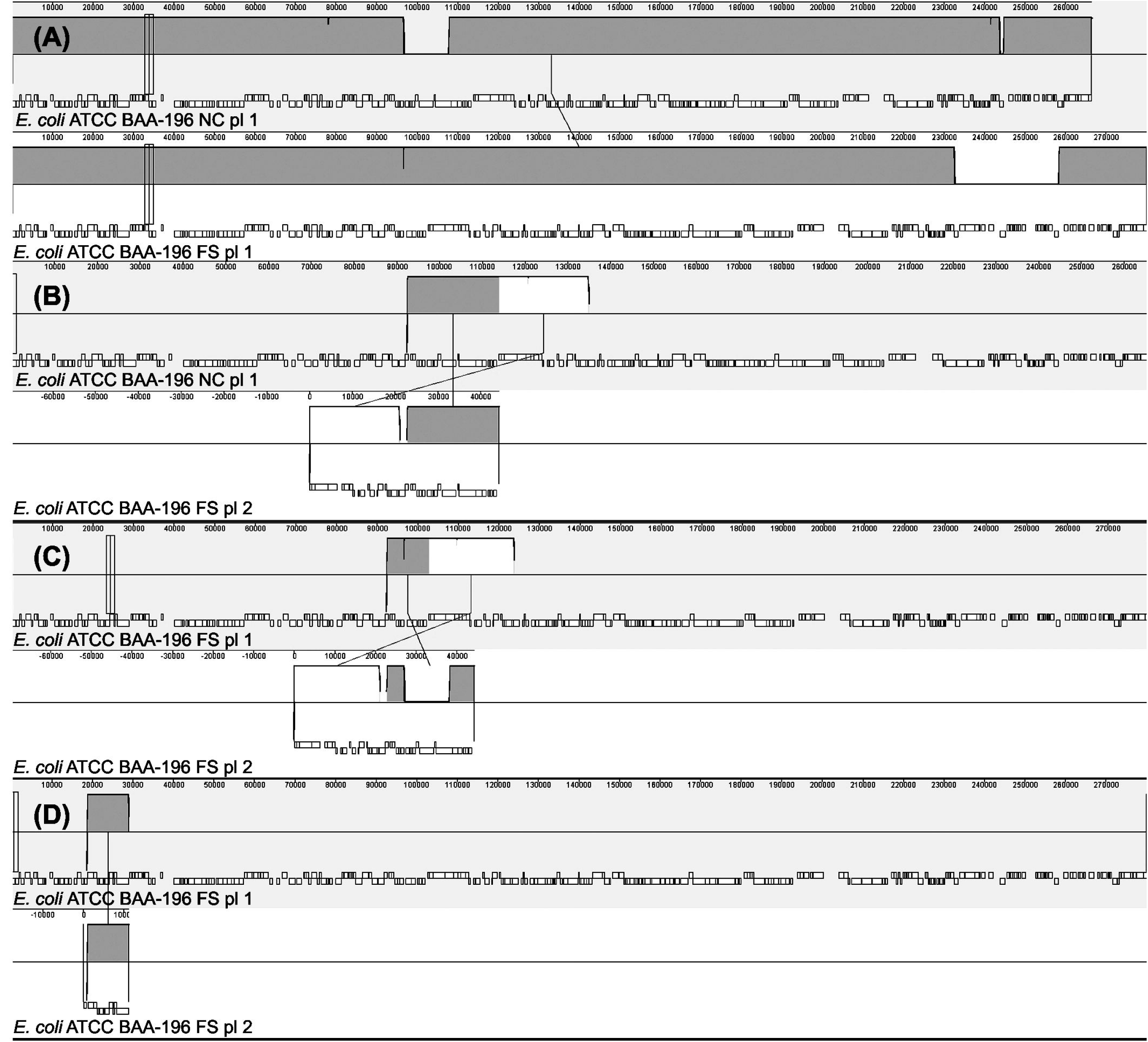
Mauve alignments of plasmid sequences of E. *coli* ATCC BAA-196. Sequence identity of homologous blocks is depicted by colored histograms. Gene locations are represented by open blocks. (A) Alignment of two large plasmids of NC and FS variants. (B) Alignment of the plasmid 2 of the variant FS against the large plasmid of the variant NC. (C) Alignment of the plasmid 2 of the variant FS against the large plasmid of the variant FS. (D) Alignment of the plasmid 3 of the variant FS against the large plasmid of the variant FS.

The identified plasmids share 90-99% sequence similarity with several known pathogenicity and drug resistance plasmids from *Klebsiella pneumonia* (pKP64477b), *Serratia marcescens*(AP014611.1) and *E. coli* (p3521). Four broad range beta-lactamases of TEM, OXA and AmpC families; five aminoglycoside inactivating enzymes of APH, ANT and AAC families; *catI* chloramphenicol acetyltransferase; trimethoprim resistant dihydrofolate reductase of the DFR family, heavy metal resistance genes *merA* and *terY*, and a tellurium resistance operon were identified on the plasmids NC and FS. Several other genes may additionally contribute to antibiotic resistance, which include 23 efflux pumps of various families and three peptide antibiotic resistance genes *bacA, pmrF* and ugd, which were predicted as potential antibiotic resistance determinants by RGI5.1.0/CARD 3.0.5 Web server.

Pathway Tools predictions showed that several genes located on the plasmid: sucrose-6F-phosphate phosphohydrolase, adenine/guanine phosphoribosyl transferase, carbamoylphosphate synthase large subunit, starvation-induced cysteine protease and citrate lyase beta subunit, may potentially interfere with several central metabolic pathways.

Two additional smaller plasmids of 44,240 bp and 11,153 bp were assembled from PacBio reads generated from *E. coli* ATCC BAA-196, variant FS. Alignment of the smaller plasmids against the large ones showed that they were generated by fragmentation and rearrangement of the large plasmid (Fig 2B-D). The middle size plasmid is constituted of two blocks linked by an integrase *retA*. This plasmid contains conjugation genes *traDGFIHJ, trhH* and some other genes associated with plasmid replication and gene integration. The smallest plasmid is a transposable element with a cluster of antibiotic resistant genes aminoglycoside 3’’-nucleotidyltransferase *aadA*, small multidrug resistance (SMR) efflux transporter *qacE*, sulfonamide-resistant dihydropteroate synthase *sull* and a GNAT family N-acetyltransferase, which are enclosed by several copies of transposases and integrases. Plasmid rearrangements and fragmentation observed in the genome of the variant FS suggests that cultivation of the strain with the sub-bactericidal concentrations of FS-1 may destabilize the initial large plasmid by activating multiple transposases and integrases found in the plasmid genome. The most likely scenario is that the large and smaller plasmids in the culture FS do not coexist in bacterial cells but resulted from fragmentation and losing parts of the plasmid in sub-populations of FS-1 treated *E. coli*.

### Differential gene regulation in *E. coli* cultivated with FS-1

Gene expression regulation in the variant FS of *E. coli* ATCC BAA-196 that survived 10 passages of cultivation with 500 μg/ml FS-1 in the medium, was compared to the NC variant cultivated for 10 passages in the same medium without the drug. The variant FS was characterized with the increased susceptibility to gentamicin and ampicillin (Fig 1) that may result from an altered gene regulation.

A volcano plot of regulated genes is shown in Fig 3. The differentially expressed genes are plotted in the central part of Fig 3 according to the fold change difference of their expression in FS culture compared to NC. Panels on the right part of Fig 3 denoted as –Inf and Inf represent respectively the genes, which were expressed only in the NC culture grown on common MH medium and the genes expressed only in FS culture grown on the medium supplemented with FS-1. Differentially expressed genes of the plasmid origin are highlighted in red.

**Fig. 3.**
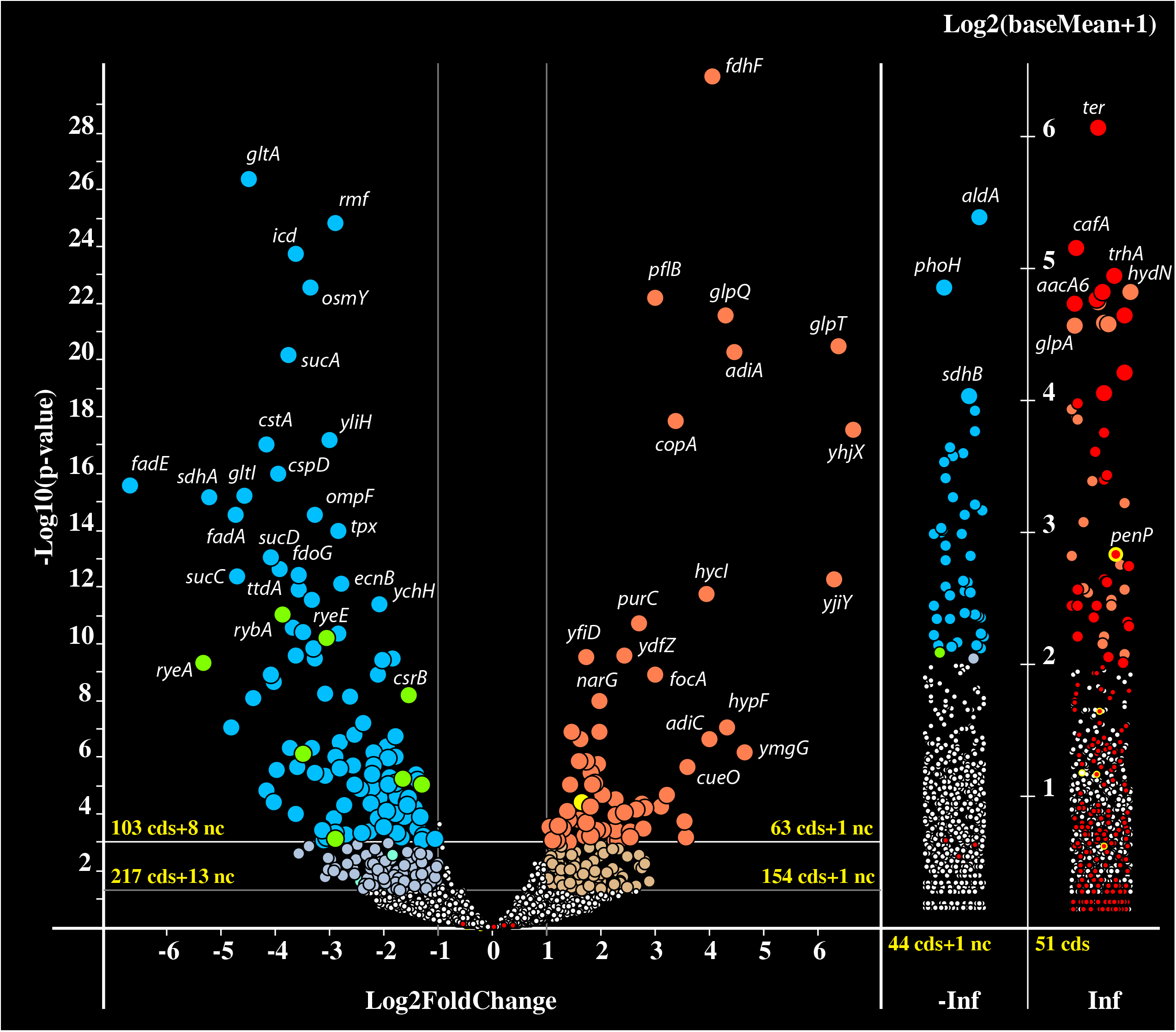
Volcano plot of gene regulation in the FS-1 treated variant (FS) of E. *coli* ATCC BAA-196 in comparison to the negative control variant (NC). Dots on the plots depict the negative (blue – CDS, green – ncRNA) and positive (red – CDS, yellow – ncRNA) gene regulation. Plasmid born genes are depicted by red circles and beta-lactamase genes located on the plasmids are highlighted by a thick yellow outline. Log_2_ fold-changes of gene expression values are on the X axes, with the common *p*-values logarithms of gene expression estimations on the Y axes. P-values were calculated by DESeq2 algorithm on three repeats of gene expression counts. Genes expressed only in the variant FS are plotted along the axis of expression counts in the positive infinity columns (Inf), and the genes expressed only in the NC variant are in the negative infinity columns (-Inf). Size of the circles and the color intensity depict estimated p-value above 0.05 and above 0.01 (bigger and brighter circles). Numbers of regulated CDS and ncRNA genes of different categories are indicated. Highly regulated genes are labeled.

The list of regulated genes is given in Supplementary Table S1. The regulated genes grouped by relevant metabolic pathways predicted by the program Pathway Tools 24.0 are shown in Table 2. Analysis of the regulated genes showed that up- and down-regulation of these genes is strongly associated with three conditions: strict anaerobic growth, low pH and osmotic/oxidative stresses.

**Table 2.**
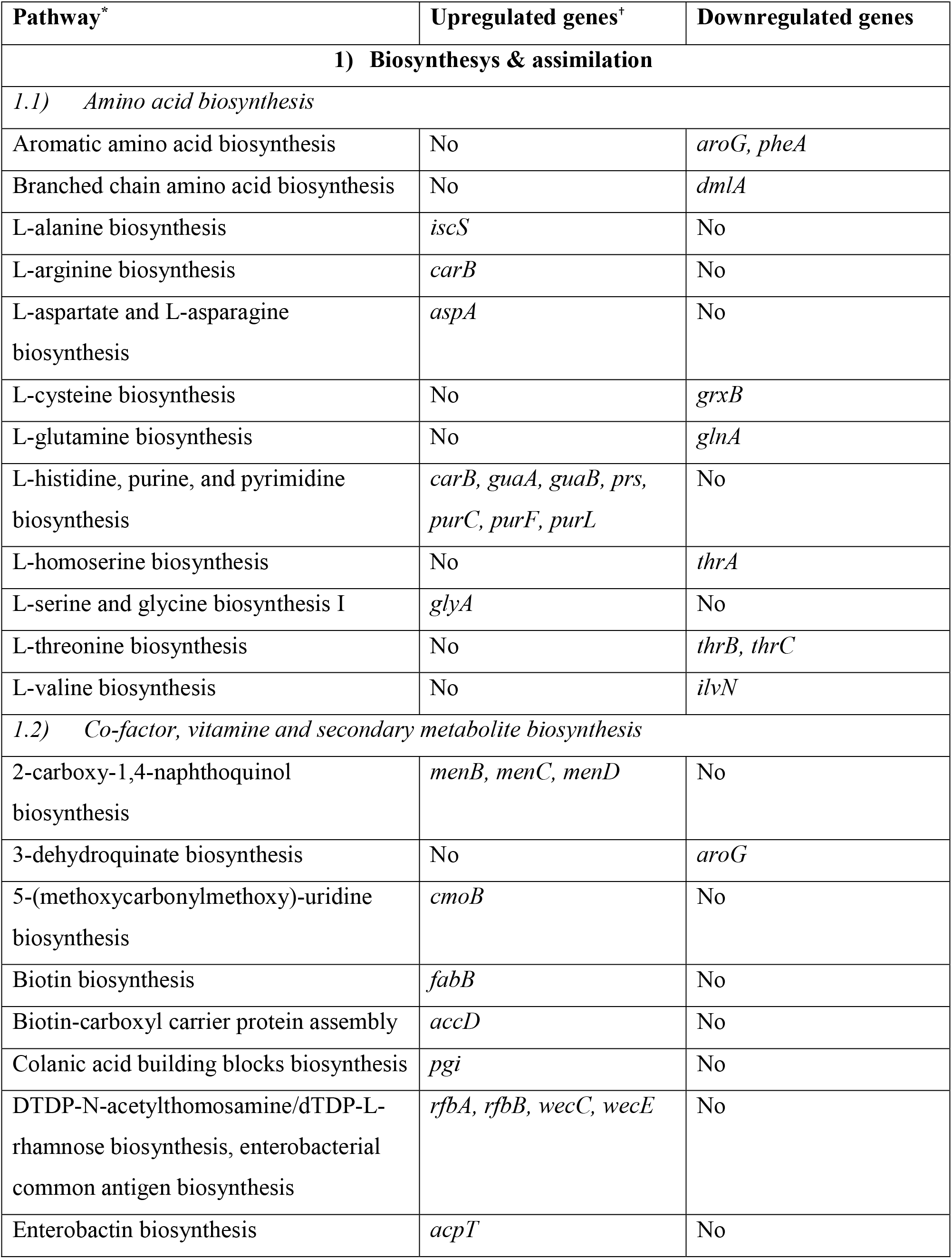
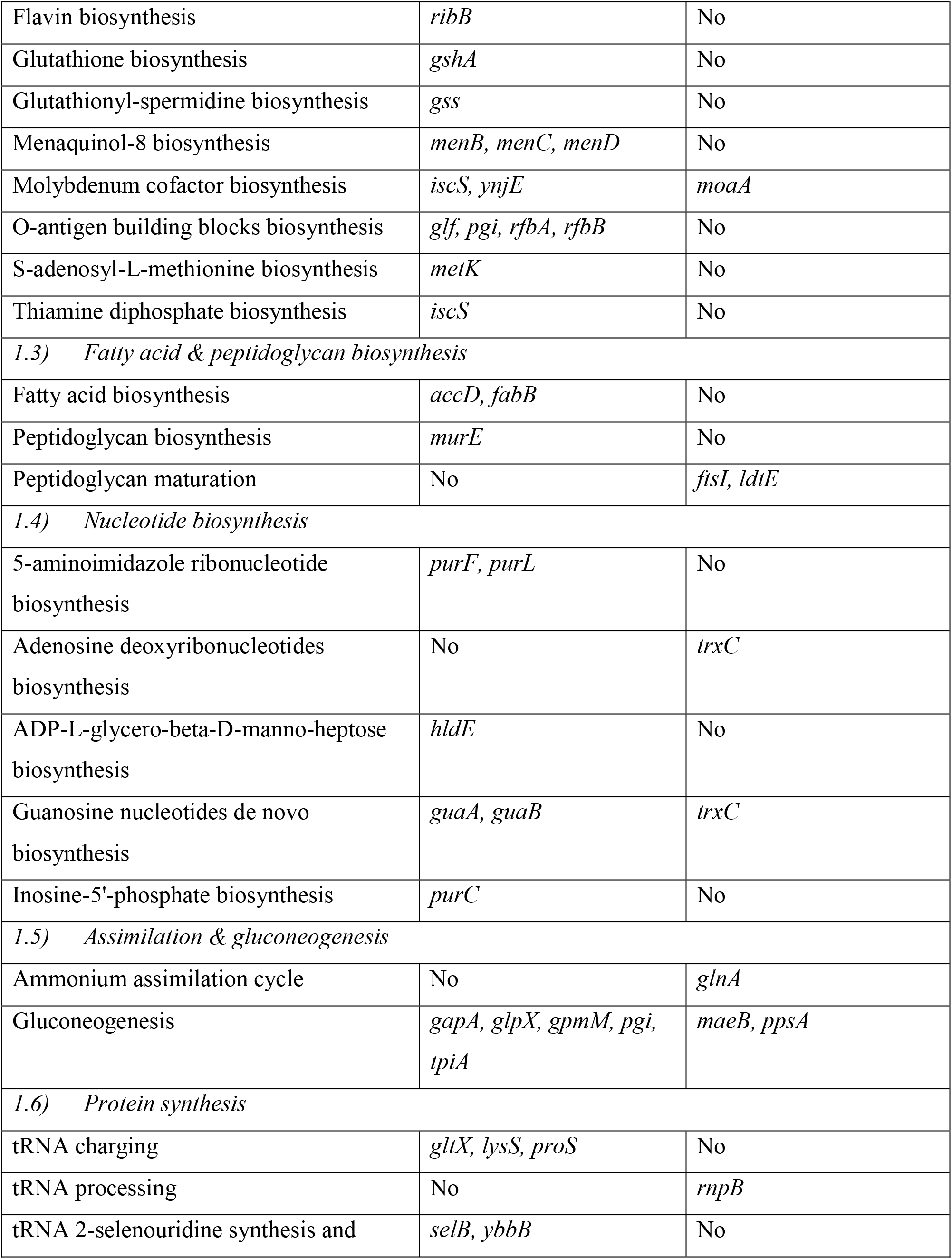
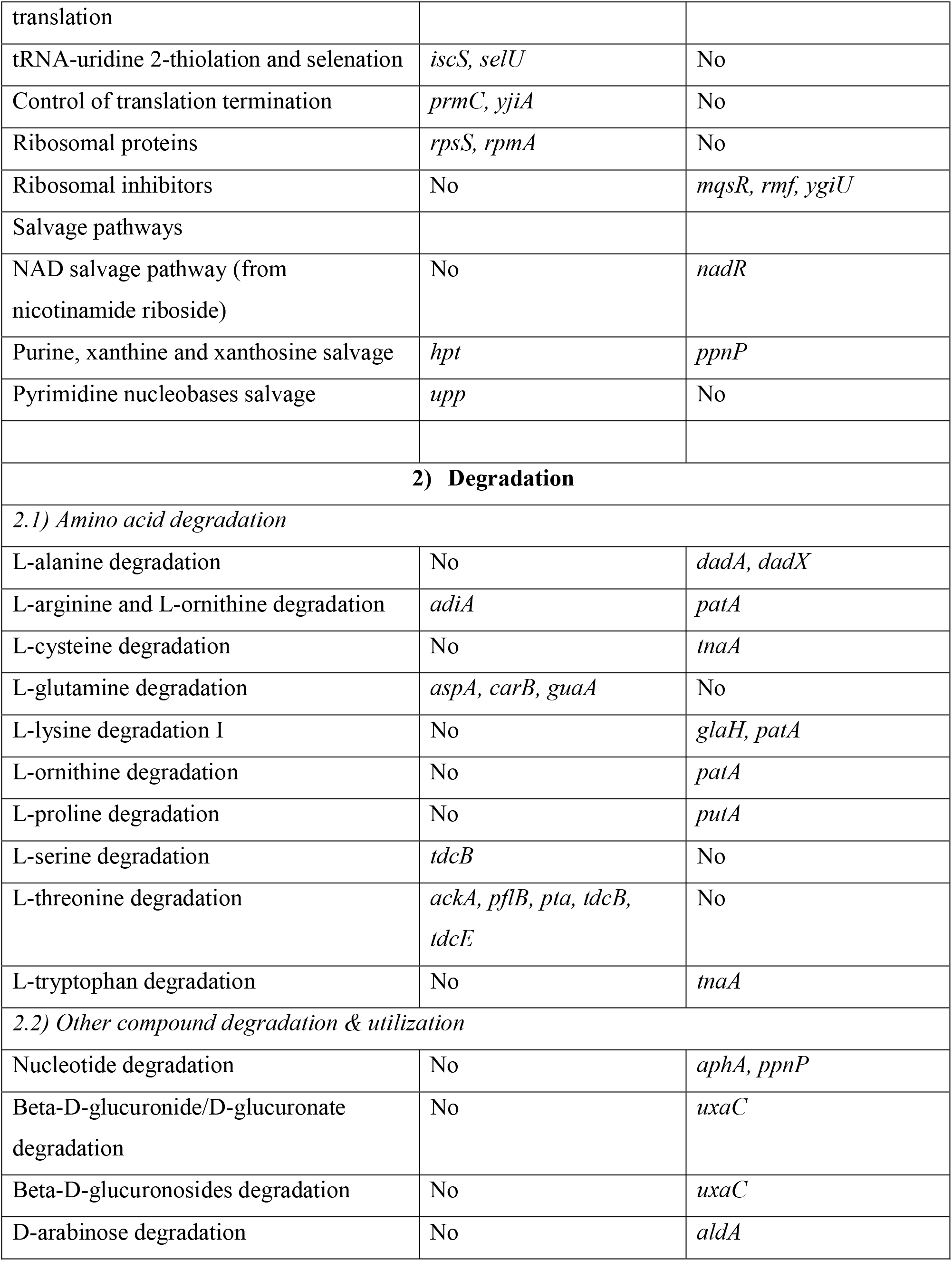
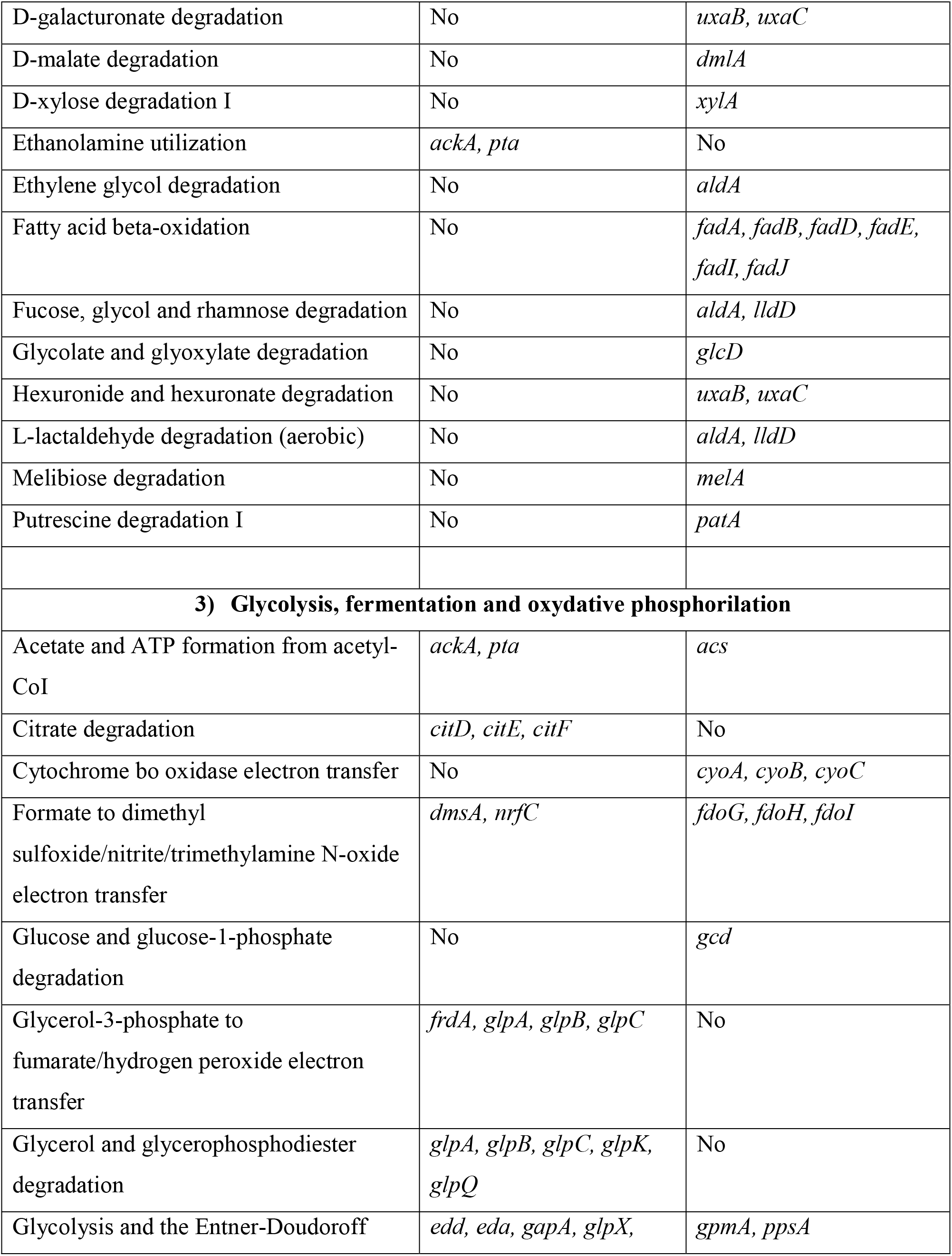
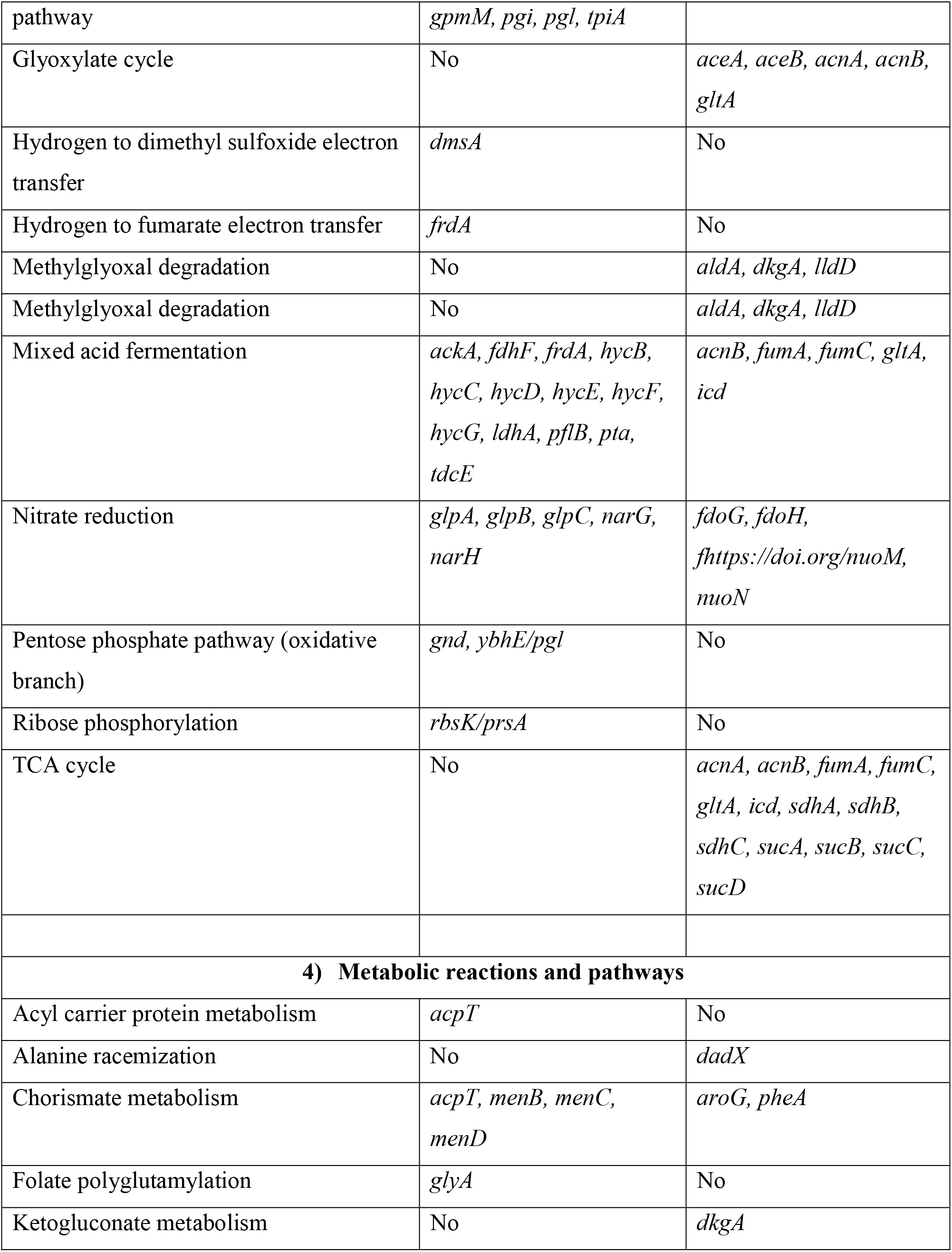
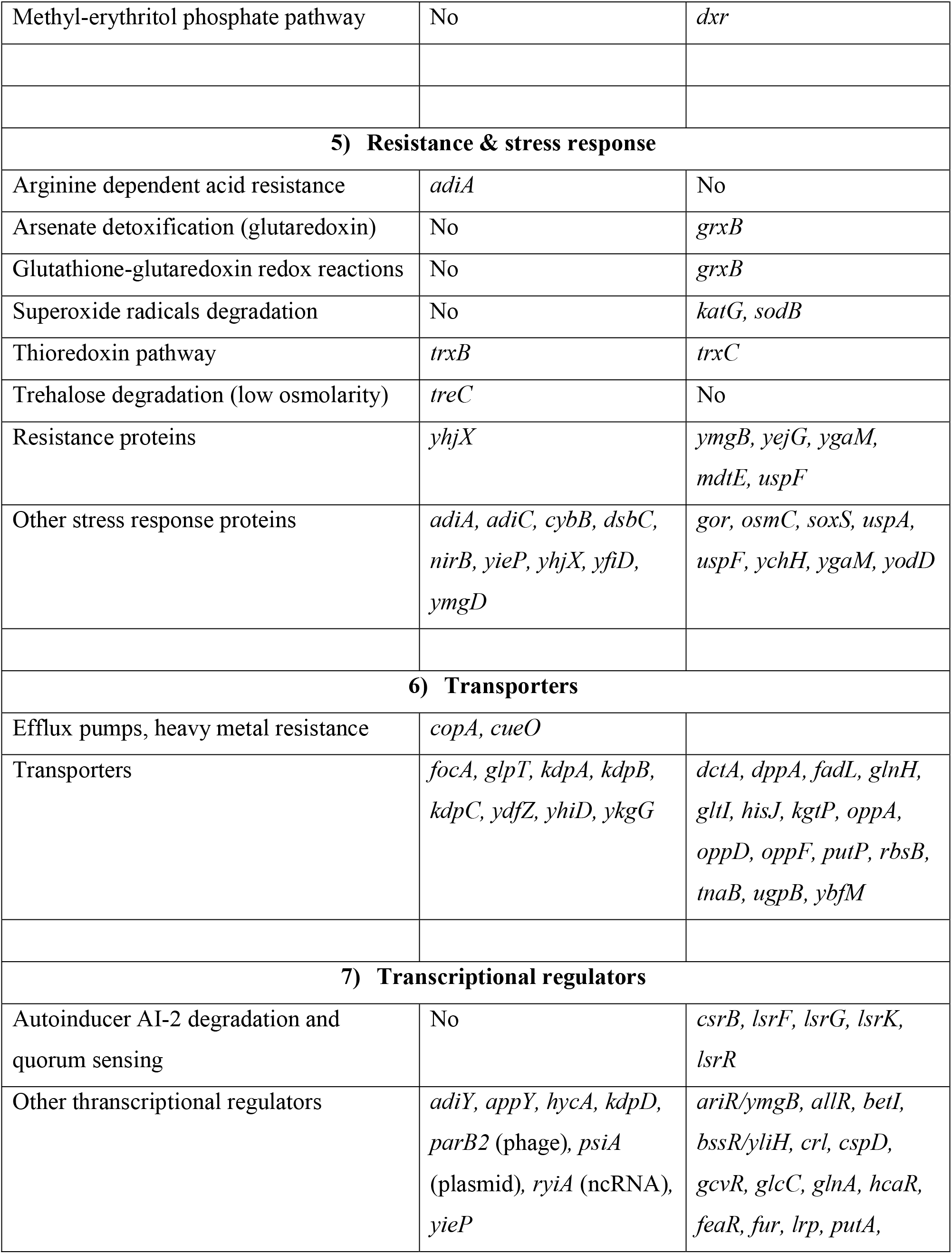
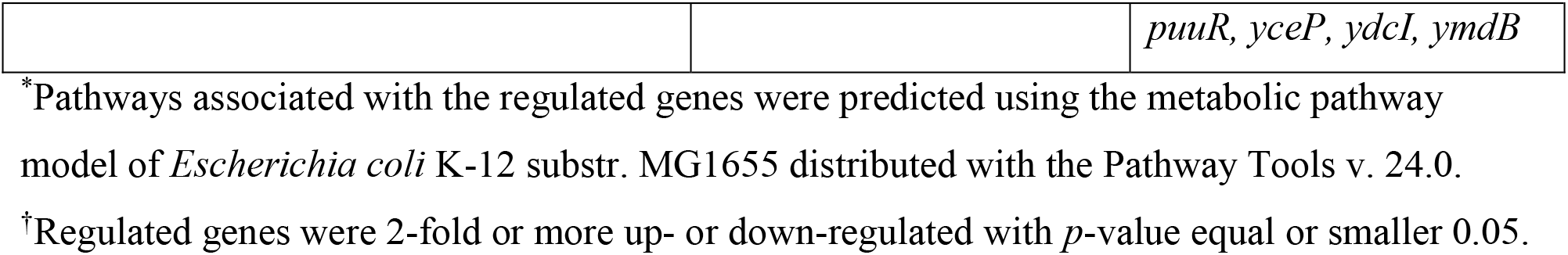
Differentially expressed genes of the culture *E. coli* ATCC BAA-196 FS compared to the culture NC and grouped by the relevant metabolic pathways.

PheNetic network of transcriptional regulation of differentially expressed genes is shown in Fig 4A. These genes were grouped into regulons controlled by the following transcriptional regulators: RpoS, Crp, ArcA, RpoH, RpoN, GcvB, InfAB and Fnr. RpoS is the master regulator of the general stress response in *E. coli*. The RpoS related stressosome is expressed in response to sudden changes in the environment to allow bacterial cells to adapt to this change [41]. The expression of this sigma-factor and controlled genes was strongly down-regulated, while the genes suppressed by *rpoS* were up-regulated indicating that the long cultivation with FS-1 required other defense mechanisms rather than the immediate stress response.

**Fig. 4.**
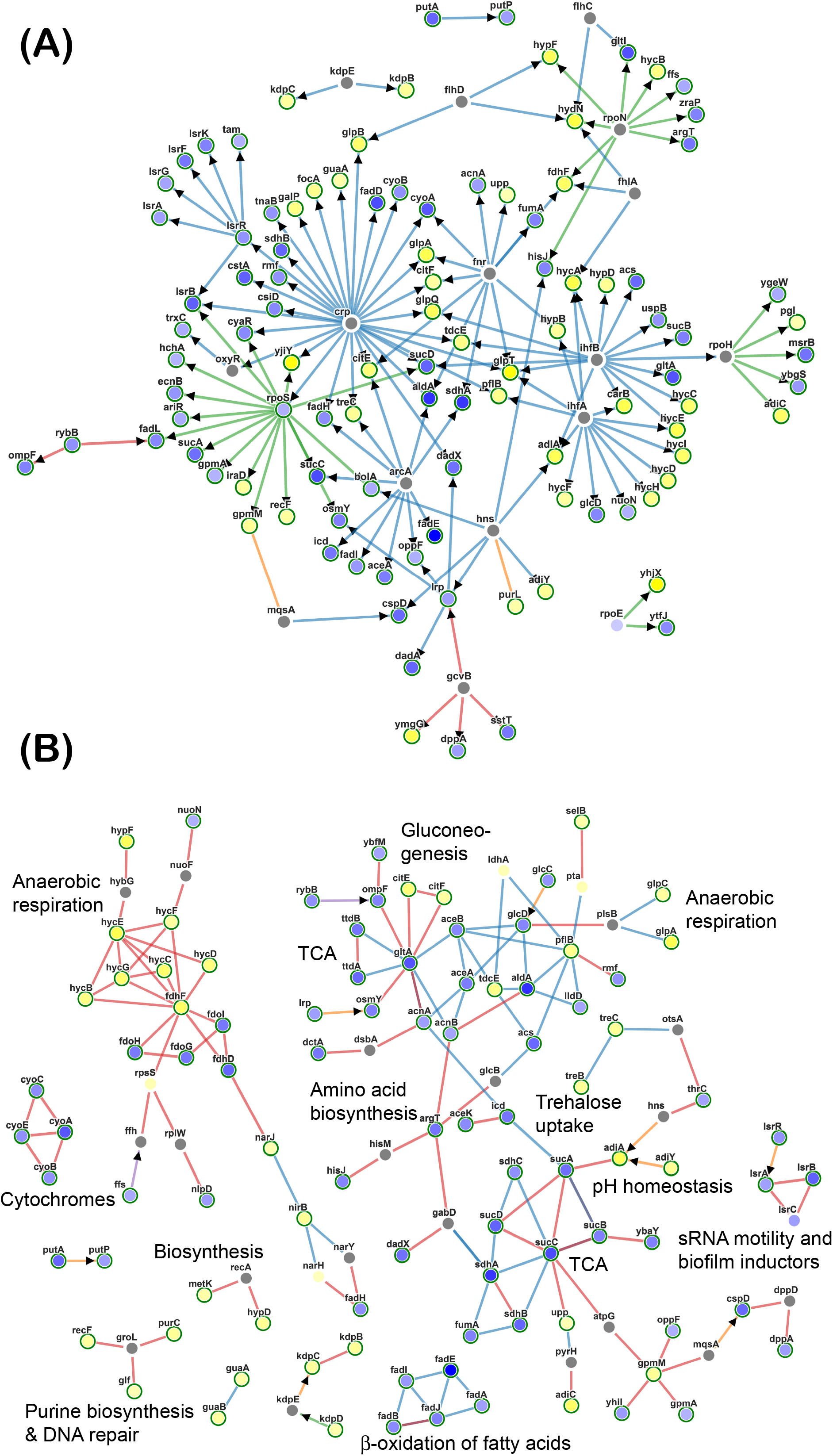
PheNetic networks of regulated genes. (A) Network of genes clustered according to the common regulation of these genes by higher level transcriptional regulators; (B) Network of regulated genes functionally link to each other. Several functional groups of genes are labeled. Upregulated genes are depicted by yellow nodes and downregulated genes – by blue nodes (vertices). Color intensity indicates the level of the regulation. Grey nodes are genes of transcriptional regulators involved in the network, which expression was not reliable changed at the aforementioned conditions. Green edges show activation relations; blue edges – inhibitory relations; and brown edges – ambivalent or neutral relations. Direct regulations by transcriptional regulators are indicated by arrowheads.

Adaptation of *E. coli* to the permanent presence of FS-1 in the medium involved a significant alteration of the central metabolic pathways (Fig 4B). An unexpected observation was that the metabolic processes were switched to anaerobic respiration dependent on formate and glycerol-3-phosphate as electron donors, and nitrates and fumarate as terminal electron acceptors. The later reactions are controlled by HydN electron carrier protein, Sn-glycerol-3-phosphate anaerobic dehydrogenase complex GlpABC, selenocysteine containing formate dehydrogenase FdhF and formate-hydrogenlyase complex HycBCDEFG, which showed no expression at the negative control condition but were highly expressed on the medium with FS-1. All systems associated with the aerobic lifestyle, such as the aerobic formate dehydrogenase complex FdoGHI, were strongly inhibited either directly by ArcA repressor or by associated regulatory pathways (Fig 4A). Expression of all genes of the cytochrome *bo* terminal oxidase complex, *cyoA, cyoB, cyoC* and *cyoE*, was strongly suppressed. It may result from damaging of cytochrome molecules by iodine released from FS-1.

An alternative hypothesis may be that the oxidative phosphorylation pathway was restrained to decrease the oxidative stress caused by the halogen. In a previous study on *Staphylococcus aureus*, the oxidative stress was reported after an exposure of the culture to FS-1 for 5 min [9, 42]. Many systems preventing cell damaging by free radicals, which are byproducts of the oxidative phosphorylation, were down-regulated in the culture cultivated with FS-1. These down-regulated genes include *tpx* thiol peroxidase; *osmC* peroxiredoxin; glutathione reductase gor; *yodD, ychH* and *ygaM* genes. Other systems probably aiding the cells to cope with halogen oxidation were activated including periplasmic disulfide oxidoreductase *dsbC* and membrane bound superoxide:ubiquinone oxidoreductase *cybB*.

All genes of the tricarboxylic acid cycle (TCA) were strongly inhibited in the FS-1 treated *E. coli* except for MaeA/SfcA NAD-dependent malate dehydrogenase that decarboxylates malate to pyruvate. However, in *E. coli* strains blocked in the fermentative pyruvate utilization, this gene is overexpressed to support cell growth by catalyzing the normally nonphysiological reductive carboxylation of pyruvate to malate [43] for further use in gluconeogenesis. Two enzymes of the gluconeogenic pathway (fructose 1,6-bisphosphatase GlpX and glucose-6-phosphate isomerise Pgi) and two enzymes shared by the gluconeogenesis and glycolysis (glyceraldehyde-3-phosphate dehydrogenase GapA and triosephosphate isomerase TpiA) were upregulated.

Anaerobic fermentation is less effective in terms of energy production compared to the oxidative phosphorylation and thus the cell requires an intensification of the glycolytic pathways and TCA cycle [44]. It contradicts to the observed inhibition of TCA enzymes and the activation of gluconeogenesis. *E. coli* has several alternative glycolytic pathways bypassing glycolysis: Entner Doudoroff (ED), Embden-Meyerhof-Parnas (EMP) and oxidative pentose phosphate (OPP) pathways [45]. The genes of the ED pathway, phosphogluconate dehydratase *edd* and keto-hydroxyglutarate-aldolase *eda*, were 1.8 and 1.6-fold upregulated. Genes *pfkA* and *pfkB* encoding subunits of the EMP pathway enzyme, 6-phosphofructokinase, were not regulated at this condition. Gnd 6-phosphogluconate dehydrogenase starting the OPP pathway was 3-fold upregulated in *E. coli* at the presence of FS-1. Upregulation was observed also for two other genes of this pathway: ribose-phosphate pyrophosphokinase *prsA* and 6-phosphogluconolactonase *ybhE*. Activation of the OPP pathway redirects the glycolytic fluxes towards the ED pathway [45] activated by FS-1. It may be concluded that the ED pathway was used by *E. coli* as the main glycolytic pathway under the effect of FS-1.

All the pathways of acetate catabolism including acetate uptake transporters and the enzymes of the fatty acid β-oxidation (aerobic and anaerobic) and the glyoxylate pathway were strongly inhibited in the FS-1 treated culture. It was unexpected as the fatty acid beta-oxidation was reported as a critical pathway during the anaerobic growth of *E. coli* on several sugars, particularly on xylose [46]. Accumulation of acetate and other organic acids produced by anaerobic fermentation and due to acetate uptake inhibition could lead to an acidic stress indicated by upregulation of several acidic stress response genes: *yhjX, ymgD, yfiD* and *adiABC*. In consistence with this was the observed upregulation of arginine decarboxylase AdiABC and lactate dehydrogenase LdhA, which are activated in anaerobic conditions in response to extremely acidic environments acidified by carbohydrate fermentation products [47, 48].

Protein and fatty acid biosynthesis pathways generally were activated in the cells treated with FS-1. Proteins PrmC and YjiA controlling the accuracy of protein translation [49, 50] and the elongation factor P *(efp)* were activated, while the ribosomal inhibitors *ygiU/mqsR* and rmf [51, 52] were strongly downregulated. At the same time, the uptake and transportation of many amino acids, organic acids, sugars and polypeptides were inhibited. Iodine in the FS-1 complex is bound to polypeptide and oligosaccharide micelles [6, 8]. Transmembrane polypeptide and amino acid transporters, such as low affinity tryptophan transporter TnaB, oligopeptide transporter OppADF, dipeptide transporter DppA, proline transporter PutP, glutamate/aspartate ABC transporter GltI and glutamine ABC transporter GlnH could serve as entry points for bound iodine atoms and thus were downregulated in the FS-1 treated *E. coli*.

Activation of heavy metal homeostasis efflux pumps *cueO* and *copA*, and potassium uptake ATPase *kdpAB* involved in osmotic stress response [53, 54] may be indicative for an increased penetrability of bacterial cell wall and cell membrane leading to leaking potassium ions from the cell and an influx of heavy metal ions from the environment. This increased penetrability of bacterial cells may be associated with damaging cell wall structures by iodine causing a destruction of protective molecular barriers. The increased penetrability of cell wall of FS-1 treated *M. tuberculosis* has been demonstrated recently [8].

Plasmid genes were transcriptionally silent when the culture was growing at the negative control condition. An exception was several transposases showing a permanent low-level expression. Cultivation with FS-1 activated most of the plasmid genes. Among the activated genes, there are four β-lactamases shown in Fig 4A. However, the highest level of activation was recorded for the genes associated with plasmid conjugation and transformation. The repressor of the cell-to-cell plasmid transfer *ybaV* [55] located on the chromosome was strongly downregulated. Hyperactivation of transposases and integrases located on the plasmid may explain the fact that two smaller plasmids were found only in the *E. coli* population grown on the medium with FS-1. Gene comparison showed that these plasmids originated from the large plasmid by genetic rearrangements (Fig 2).

Despite of the activation of the plasmid born β-lactamases, viability of FS-1 treated *E. coli* decreased and susceptibility to antibiotics increased (Fig 1). It may result from the observed downregulation of many other genes associated with drug resistance: *yejG* [56], *uspF* [57] and *ygaM* [58], and also with the oxidative stress and increased penetrability of the cell membrane. Strong inhibition of the genes involved in biofilm formation was observed in the FS-1 treated culture that potentially can reduce pathogenicity of the bacterium. Expression of quorum sensing regulators *lsrR* and *csrB* [59-61] was strongly downregulated. Directly or indirectly, the inhibition of regulatory elements has affected many other genes involved in bacterial motility, adhesion and biofilm formation, which include the AI-2 quorum sensing signal procession protein [52], AriR transcriptional regulator [62], inducer of biofilm formation BolA [63], stress induced protein *uspF* [57], biofilm-stress-motility lipoprotein BsmA/YjfO [64] and the regulator of biofilm formation through signal secretion protein BssS/YseP [65].

It should be noted that the increased susceptibility of FS-1 treated *E. coli* to gentamicin and ampicillin (Fig 1) was observed after washing the bacterial cells from the drug FS-1 when the direct antibacterial effect of the drug was substantially decreased. It may be supposed that cultivation of *E. coli* with FS-1 caused profound epistatic and/or epigenetic adaptive changes in the bacterial population affecting its susceptibility to other antibiotics, which may persist in several generations. To investigate this hypothesis, epigenetic modification in chromosomal DNA of FS-1 treated *E. coli* were studied.

### Epigenetic modifications of chromosomal nucleotides

Epigenetic modifications of nucleotides of chromosomal and plasmid sequences of NC and FS variants of *E. coli* ATCC BAA-196 were predicted by the kinetic analysis of base calling during SMRT PacBio sequencing. The statistical reliability of epigenetic modifications was calculated as Phred-like quality base modification (BM) scores. The values above 21 correspond to the confidence *p*-value 0.01 or smaller. PacBio sequencing of the control and FS-treated bacterial samples was performed in tree repetitions followed by calculation of average BM scores for every nucleotide at both strands of the genome. Rank diagrams of distribution of BM scores in NC and FS genomes are shown in Fig 5. SMRT Link program predicted that the peaks of high scored adenine residues in both genomes (BM scores > 80) pertained to methylation of the chromosomal DNA at these sites.

**Fig. 5.**
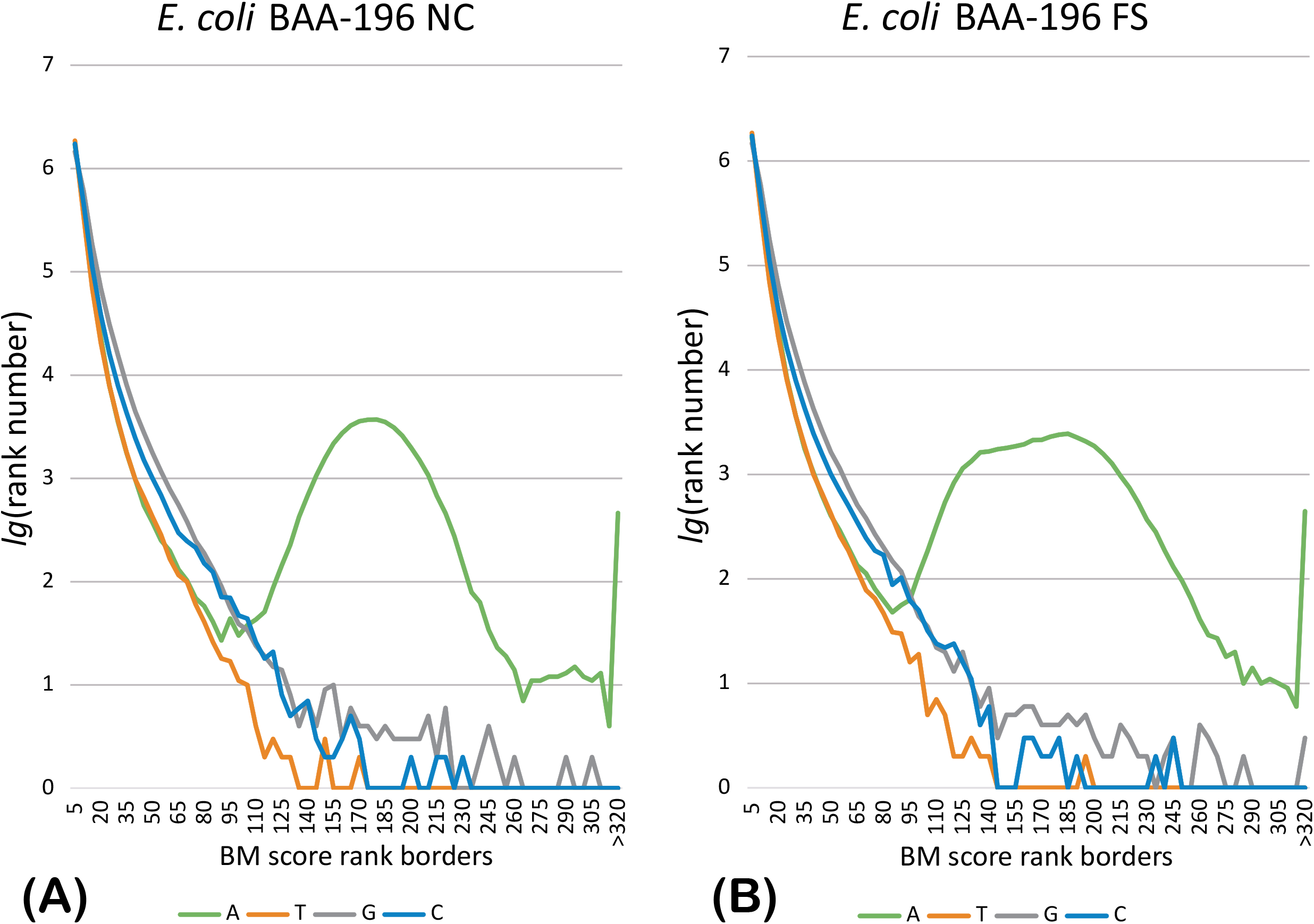
Distribution of BM scores of modified nucleotides. Plot (A) shows the distribution in the negative control (NC) variant of *E. coli* ATCC BAA-196; plot (B) shows the distribution in the FS-1 treated (FS) variant. Axis X depicts score ranks stepping 5 score units. Axis Y indicates decimal logarithms of rank numbers. BM scores in the graphs are average values calculated in three repeats for every individual set of NC and FS DNA reads generated from repeated DNA samples.

Methylation motifs and types of methylations were predicted by the program *motifMaker*.Frequencies of different methylation motifs in the sequenced genomes are shown in Table 3. Three adenine methylation motifs were identified: more abundant type-II methylation G**A**TC, and two motifs of type-I methylation A**A**C(N_6_)GTGC and GC**A**C(N_6_)GTT, which to some extend are reverse complement sequence of each other. (Methylated nucleotides in the motif sequences are shown in bold and, in the case of a double strand methylation, the nucleotides opposing the methylated nucleotides on the reverse complement strand are shown in italic.) G**A**TC and A**A**C(N_6_)GTGC / GC**A**C(N_6_)GTT methylation was exhaustive in the sequenced genomes covering 100% of all available sites; however, only 73-76% of the available motifs were methylated at both DNA strands (Table 3). Bipartite adenine methylation in sequence motifs G<u>A</u>*T*C controlled by S-adenosylmethionine depended orphan (not associated with any restriction enzymes) Dam methylase is common in *E. coli* [66, 67]. G**A***T*C sites were often associated with a more complex pattern of nucleotide modifications, **C**RGKG**A***T*C, where the leading cytosine residue was also methylated. While this type of cytosine methylation was not the most abundant, these sites of modified cytosines showed the highest BM scores. Notably, while the total numbers of **C**RGKG**A***T*C sites in NC and FS genomes remain similar (respectively 105 and 106 out of 745 available CRGKGATC motifs), they were found at different loci on the chromosomes (Fig 6A-B).

**Table 3.** Methylation motifs predicted by the program *motifMaker* in three repeats of SMRT PacBio sequencing of the complete genome of NC and FS variants of *E. coli* ATCC BAA-196.

| Methylation motifs | NC variant |  | FS variant |  |
| --- | --- | --- | --- | --- |
|  | Chromosome | Plasmid | Chromosome | Plasmid |
| GATC* | 28426 / 38442 (74%)<sup>†</sup> | 1151 / 1854 (62%) | 28408 / 38442 (74%) | 1218 / 1950 (62%) |
| AAC(N6)<u>GTGC</u> | 437 / 602 (73%) | 14 / 20 (70%) | 437 / 602 (73%) | 16 / 21 (76%) |
| GCAC(N6)<u>GTT</u> | 447 / 602 (74%) | 13 / 20 (65%) | 447 / 602 (74%) | 14 / 21 (66%) |
| CCAGG | 622 / 12113 (5%) | 28 / 566 (5%) | 651 / 12113 (5%) | 30 / 596 (5%) |
| <u>CRGKGATC</u> | 105 / 745 (14%) | 5 / 39 (13%) | 106 / 745 (14%) | 3 / 42 (7%) |
| WCCCTGGYR | 109 / 407 (27%) | 3 / 21 (14%) | 103 / 407 (25%) | 1 / 36 (3%) |

\*In the sequences of motifs, the methylated nucleotide is underlined and, in the case of a double strand methylation, the nucleotide opposing the methylated nucleotide on the reverse complement strand is shown in italic typeface;
†Numbers of methylated sites is displayed in the following format:
*methylated\_motif\_number / total\_number\_of\_motif\_sequences (fraction of methylated motifs)*. It should be noted that only those methylation sites were counted, which were predicted in all three repeats of genome sequencing. This approach improved the statistical reliability of the positive predictions; however, it could lead to an increase of false-negative predictions.

**Fig. 6.**
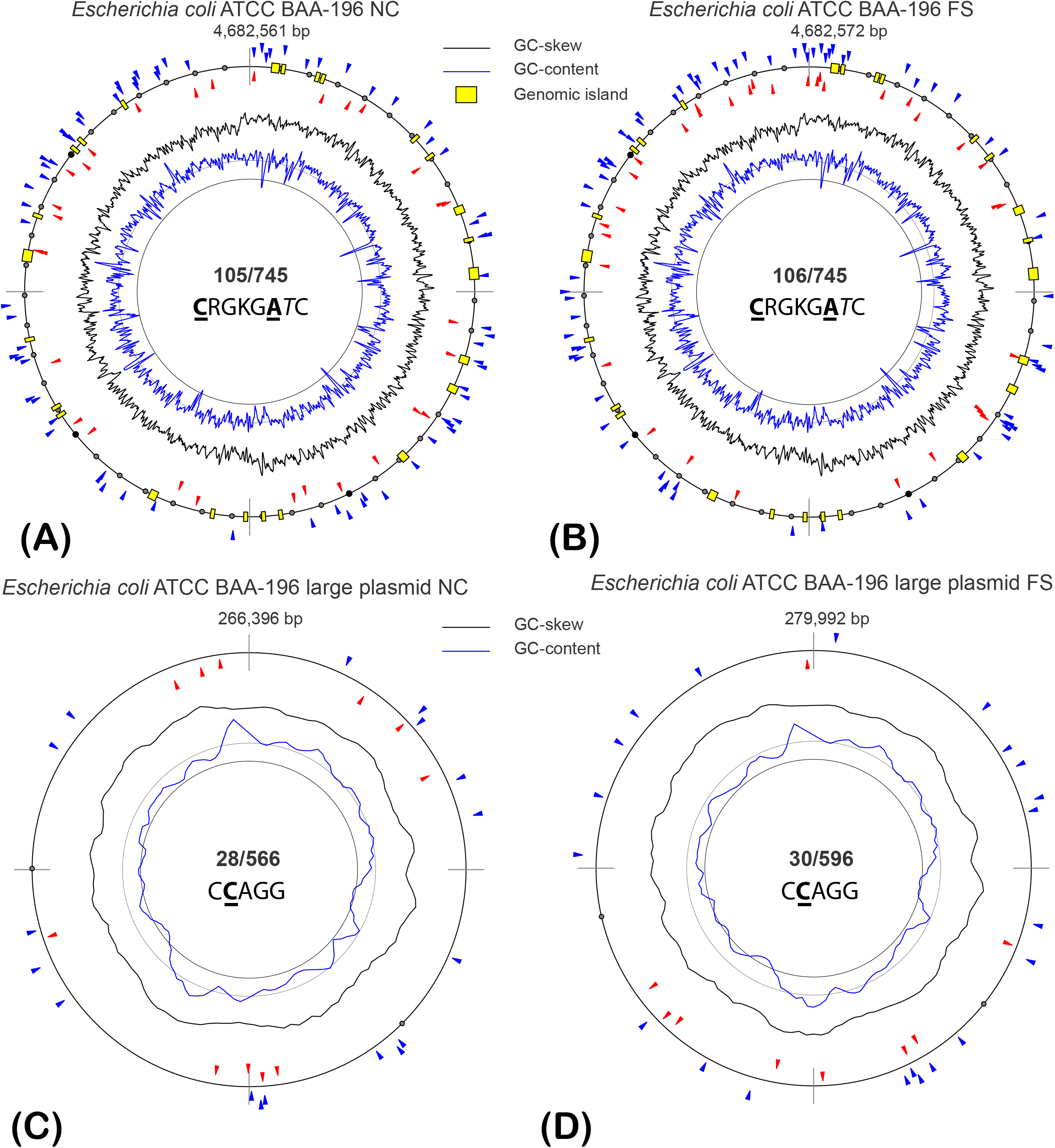
Alternative cytosine methylation. Patterns associated with **C**RGKG**A***T*C motifs in the chromosomes (A-B) and C**C**AGG motifs in the large plasmids (C-D) of the NC and FS variants of *E. coli* ATCC BAA-196 are shown. Triangle marks indicate locations of methylated sites on the direct (blue marks) and complement (red marks) DNA strands. Only those sites were selected for the analysis, which showed statistically reliable epigenetic modification predictions (BM score ≥ 21) in all three repeats. Fluctuations of GC-content and GC-skew parameters calculated for 8 kbp sliding window stepping 2 kbp are shown by corresponding histograms. Locations of identified genomic islands in bacterial chromosomes are indicated by yellow blocks.

Another type of bipartite methylation at palindromic motifs A**A**C(N6)GTGC and GC**A**C(N6)G*T*T is under control of type-1 *Eco*KI-like methylase [68]. In the sequenced genomes, the most likely candidate for this methylase is chromosomal gene *hsdM* within an operon comprising also type I restriction enzyme *hsdR* and specificity determinant *hsdS*. This operon is followed by three other restriction-modification genes, uncharacterized gene *yjiV* and two subunits of 5-methylcytosine-specific restriction enzyme McrBC, which were upregulated in the culture FS (78-fold for *mcrB*), while the operon *hsdRMS* was constantly expressed in both cultures. Methylation at motifs A**A**C(N_6_)G*T*GC / GC**A**C(N_6_)G*T*T protects DNA from cleavage by the cognate restriction enzyme. However, approximately 25% of these palindromic motifs were methylated only at one DNA strand (Table 3). Either one strand methylation was sufficient to protect DNA from cleavage, or it could be that the semi-palindromic nature of these motifs excludes its recognition by protein HsdS on both DNA strands, or these sites could be protected from both, methylation and restriction, by some regulatory elements (DNA binding proteins or small RNA molecules). Also, the stringent selection of only those modified sites which were predicted in all three repeats of the experiment potentially could increase the rate of falsenegative predictions.

Other predicted cytosine methylation motifs were C**C**AGGRAH and W**C**CCTGGYR. Methylation at C**C**AGG Dcm motif is typical for *E. coli* [66]; however, *motifMaker* has predicted in NC and FS genomes a similar but more complex methylation motif C**C**AGGRAH. Only a fraction of methylated sites C**C**AGG was associated with 3’-end sequence conservation; however, it may explain why this palindromic sequence was methylated only at one DNA strand. Only 5% of available CCAGG sites were methylated on the chromosomes and plasmids. While the number was constant, different sites were methylated in the NC and FS genomes. The distribution of methylated C**C**AGG motifs in the plasmid sequences of two genomes is shown in Fig 6C-D. In sequences WCCCTGGYR, despite of the obvious palindromic nature of this motif, bipartite methylation at both DNA strands took place only in 10% of the methylated sites and was asymmetric: W**C**CCTGGYR. Interestingly, this type of methylation was more common for the chromosomal rather than for the plasmid DNA. Fractions of methylation of all other motifs were similar on the chromosomes and the plasmids (Table 3).

CCAGGRAH and WCCCTGGYR sequences share the same central motif CCWGG. Nevertheless, they show different patterns of distribution and are likely controlled by different methyltransferases. Methylation of CCWGG motifs is controlled by Dcm cytosine methylase also referred to as Mec methylase [69]. Two alleles of *dcm* genes associated with EcoRII-like restriction-modification systems were found in the genome of *E. coli* ATCC BAA-196 with chromosomal and plasmid locations: BAA196NC_1741 and BAA196NC_RS24085, respectively. Both these genes are expressed at NC and FS conditions (plasmid gene showed 4fold transcriptional activation; however, it was not statistically reliable). Chromosomal gene *dcm* is followed on the chromosome by short patch repair DNA mismatch endonuclease *var*. It shows that this methylation may be associated with DNA repair mechanisms rather than with DNA cleavage prevention [67]. Contrary, DNA cytosine methyltransferase located on the plasmid is neighbored with a type II restriction endonuclease BAA196NC_RS24090.

Only a small fraction of CCWGG motifs were methylated. It may be explained by the fact that *Eco*RII restriction enzymes do not cleave DNA at a single recognition site but require binding to an additional target site serving as an allosteric effector [70]. Thus, restriction of CCWGG sites in *E. coli* may depend on the spatial conformation of the chromosomal and plasmid chromatin, which may be guided by methylation of CCWGG sites.

On the chromosomes NC and FS, the total numbers of adenine residues with BM scores > 21 including sporadic sites not associated with the recognized motifs were 43,361 and 43,760, respectively. Among them, 41,352 sites (95%) were located exactly at the same positions in both genomes. The numbers of modified adenine residues with BM scores > 80 were 39,098 in NC and 39,083 in FS, and 39,045 (99%) were the same loci in both genomes. Only 53 high scored modified adenine residues were found in NC and 38 in FS. NC-specific adenine methylation sites were found within sequences of 16 genes, but only one of them was significantly downregulated – *yebV* encoding an uncharacterized protein. Seven genes in the genome FS contained genome-specific high scored methylated adenine residues. One of these genes, nitrite reductase *nitB*, was significantly upregulated.

There were 10,915 methylated cytosine residues in the genome NC and 11,220 in the genome FS with only 6,001 methylated sites (54%) shared by these two genomes indicating that cytosine methylation patterns were more dependent on the growth conditions. Numbers of modified cytosine residues with BM scores > 80 were 201 and 203 in NC and FS genomes, respectively; among which 119 sites (59%) were common for both genomes. NC-specific modified cytosine residues were found in 33 genes. Two of these genes, citrate-ACP transferase *citF* and DNA-binding transcriptional activator *gadW*, were significantly upregulated. One **C**RGKG**A***T*C site with NC-specific cytosine methylation was near the promoter region of the NADP-dependent malate dehydrogenase *maeB*, which was strongly downregulated by the presence of FS-1 in the medium. In contrast to FS-1 activated NAD-dependent malate dehydrogenase MaeA, MaeB activity is associated with acetate metabolism [71] that generally was downregulated in the FS-1 treated *E. coli*.

FS-specific cytosine methylation was found in 39 genes with three of them upregulated: biodegradative arginine decarboxylase *adiA*, DNA-directed RNA polymerase *rpoC* and glucose-6-phosphate isomerase *pgi*; and two genes strongly downregulated: bifunctional isocitrate dehydrogenase *aceK* and acyl-CoA dehydrogenase *fadE*.

### Complexation of FS-1 with bacterial DNA

Epigenetic changes in bacterial chromosomes under the effect of the iodine-containing nano-molecular complex FS-1 suggested a possible complexation of DNA with nanoparticles and/or with released iodine, which potentially can halogenate chromosomal nucleotides. Iodine may halogenate DNA nucleotides mostly at thymine residues [72], however, other computational simulations showed that FS-1 micelles may interact with purine residues [8]. The number of thymine residues with BM scores > 21 increased insignificantly from 3,199 sites in NC to 3,306 in FS. To study the ability of FS-1 to reach the chromosome and create complexes with DNA, the drug was synthesized with the radioactive isotope ^131^I (20 MBq/ml). After 1 h cultivation with the radioactive labeled FS-1, bacterial cells were washed twice to remove the remaining micelles of FS-1 and the residual radioactivity was measured. Then the residual radioactivity was measured in DNA samples extracted from the washed cells. It was found that radioactivity of the extracted DNA was 43.46 ± 13.895 Bq/ng that constituted 0.46% ± 0.15 of the residual radioactivity of the treated bacterial cells after washing. This insignificant residual radioactivity could be associated with either complexing of FS-1 with the DNA or a direct halogenation of nucleotides by iodine isotopes that potentially can damage DNA.

The analysis of expression of the genes responsible for DNA repair showed that the genes of the SOS-response, *recA* and *lexA*, were downregulated. Strong upregulation was detected for the gene *yjiA*, which is expressed in response to mitomycin C treatment causing DNA damage [73]. Genes *iraD/yjiD* encoding an inhibitor of the σ^S^ activity in response to DNA damage [74] also were upregulated. Genes of the DNA repair RecFOR complex were differentially regulated. RecF was strongly upregulated while RecO and RecR were downregulated. RecFOR independent double-strand break repair protein YegP was downregulated. It indicates that damaging of the chromosomal DNA of *E. coli* most likely was not the major target of the drug FS-1 and the DNA of this bacterium was protected from the direct iodination. The activated DNA repair genes may be responsible for the DNA protection from iodine.

## Discussion

The iodine-containing nano-micelle complex FS-1 was developed as a new supplementary drug against MDR-TB [10, 11]. The efficacy of the drug was confirmed in three rounds of clinical evaluation of FS-1 (www.clinicaltrials.gov, acc. NCT02607449). Studying of mechanisms of the therapeutic activity of this drug was hampered by obvious technical problems of using MDR-TB isolates as model organisms. Reproduction of antibiotic resistance reversion by FS-1 on *E. coli* [12] and *S. aureus* [9, 13] facilitated further studies and demonstrated opportunities of designing of new iodine-containing nano-micelle drugs against multidrug resistant nosocomial infections. An experimental scheme to study the long-lasting effect of FS-1 on antibiotic susceptibility of treated microorganisms first was applied on *S. aureus* ATCC BAA-39 [9] and currently on *E. coli* ATCC BAA-196 that allows comparison of effects of the drug on representatives of Gram-positive and Gram-negative pathogenic microflora.

*E. coli* ATCC BAA-196 is a laboratory strain constructed by inserting plasmid pMG223 from *K. pneumoniae* ATCC 14714 bearing multiple antibiotic resistance determinants into nonpathogenic and antibiotic-susceptible strain *E. coli* J53-2 [14]. Contrary, the strain *S. aureus* BAA-39 used in other study is a natural MRSA strain bearing a horizontally acquired methicillin resistance cassette SCCmec [13].

No plasmid gene expression was observed in *E. coli* BAA-196 on the regular medium without antibiotics. Plasmid gene expression was stimulated by the presence of FS-1; however, the susceptibility of the treated strain to gentamicin and ampicillin has increased (Fig 1). This observed positive therapeutic effect of the treatment may be explained, at least partly, by a destabilizing effect of FS-1 on acquired virulence plasmids and genomic islands. This study showed that in the treated *E. coli* population, there was a tendency towards plasmid fragmentation due to activation of plasmid-born transposable elements by abiotic stresses associated with the FS-1 treatment. A similar destabilizing effect of FS-1 leading to an increased rate of excision of SCCmec cassettes was reported on *S. aureus* [9]. Loss of plasmid located virulence factors and plasmid rearrangements under stressful growth conditions (low temperature) was reported by other authors for *Aeromonas salmonicida* [75].

Another important mechanism of the action of FS-1 may consist in increasing penetrability of the cell wall and cytoplasmic membrane due to destruction of their barrier functions by iodine. First the increased permeability of cell membranes of bacteria treated with FS-1 was demonstrated on *M. tuberculosis* [8]. This study showed that *E. coli* cells shortly treated with FS-1 permeated a higher concentration of intracellular antibiotic that would obviously lead to a higher susceptibility of this culture (Table 1). However, this mechanism may explain only the direct effect of FS-1 on bacterial cells but it cannot explain the residual increase of antibiotic susceptibility of the FS-1 treated bacteria after washing the drug out.

In both model organisms, *E. coli* and *S. aureus*, the strongest activation of gene expression was reported for heavy metal efflux pump *copA*; osmotic stress response K^+^-transporters *kdpB, kdpC* and *kdpE*; and carbamoyl-phosphate synthetase subunit *carB* that possibly might participate in the stringent response [76]. Therefore, both cultures suffered from similar stresses; however, it was evident that *S. aureus* was more vulnerable to the effect of FS-1 than *E. coli*. The calculated MBC for *E. coli* BAA-196 (1,000 μg/ml) was higher than for *S. aureus* BAA-39 – 900 μg/ml. The residual radioactivity of DNA samples extracted from *E. coli* cells treated with FS-1 marked by ^131^I isotope was only 0.46% ± 0.15, whereas the experiment carried out on *S. aureus* showed 12.76% ± 9.04 residual radioactivity of DNA samples [9]. This substantial difference in in iodine permeability of cell walls of Gram-positive and Gram-negative bacteria may explain several other differences in the effect of FS-1 on *E. coli* and *S. aureus*.

Complete genome sequencing of the negative control (NC) and FS-1 treated (FS) cultures of *E. coli* BAA-196 did not reveal any significant mutations in these genomes to explain the difference in susceptibility to antibiotics. Contrary, multiple frame-shift mutations were accumulated in the FS culture of *S. aureus* BAA-39 compared to NC culture of this strain [9]. It was hypothesised that in *S. aureus*, iodine released from FS-1 could halogenate or oxidizes DNA nucleotides directly. This hypothesis was confirmed by the increased number of epigenetically modified nucleotides in the FS-1 treated *S. aureus* randomly distributed on the chromosome and not associated with any recognition motifs [9]. Such the effect of FS-1 was not observed on *E. coli*.

The antibiotic susceptibility tests were performed in the medium containing no FS-1 to minimize the direct synergetic effect of the drugs with antibiotics. This deferred effect of FS-1 on the antibiotic susceptibility of the treated culture may be explained by long-lasting genetic, epistatic and/or epigenetic changes in the treated bacterial populations. Gene expression comparison showed significant changes in the metabolism of bacteria treated with FS-1. *E. coli* cultivated with FS-1 has switched to anaerobic respiration with strong inhibition of TCA genes but activated gluconeogenesis. It was hypothesized that these metabolic alterations were to cope with the oxidative stress and the need for repairing the damaged cell wall and cytoplasmic membrane structures. In association with a strong inhibition of many transmembrane peptide and amino acid transporters, strong activation of the bacterial cell anabolism could lead to nutrient stringency and weakening of cell defense and antibiotic resistance systems.

This gene regulation observed in the FS-1 treated *E. coli* was to that observed in *S. aureus* except for an activation of glycolysis instead of gluconeogenesis. During anaerobiosis, *S. aureus* cannot help but use the glycolysis to feed the activated fermentation pathways as this bacterium, in contrast to *E. coli*, has no alternative glycolytic pathways such as the Entner Doudoroff pathway that was activated in the FS-1 treated *E. coli*. This may be another factor making *S. aureus* more susceptible to FS-1. Another noteworthy difference in gene regulations by FS-1 in *S. aureus* compared to *E. coli* was a strong activation in the former organism of multiple chaperons and DNA repair genes that was not the case with *E. coli* [9]. Contrary, *S. aureus* genes encoding cytochromes were not as much inhibited as in the FS-1 treated *E. coli*.

Long-lasting effects on gene regulation may be associated with different patterns of methylation of DNA nucleotides. *E. coli* in contrast to *S. aureus* comprised a bigger number of methyltransferases catalyzing the methylation of both adenosine and cytosine residues.

This study showed that the treatment of *E. coli* with FS-1 altered the pattern of methylation of the chromosomal and plasmid DNA that in the case with *S. aureus* was observed to a smaller extent [9]. In the strain *E*. coli BAA-196, the dominant type of DNA modification was adenine methylation at G**A***T*C motifs recognized by an orphan Dam methylase. While this methylation usually covers almost 100% available motifs, demethylation of a single nucleotide at one DNA strand may cause a significant impact on gene regulation and bacterial virulence [56, 57]. In this work, it was shown for the first time that the abundant in gamma-Proteobacteria G**A***T*C Dam methylation was associated in this bacterium with a more complex pattern of tripartite cytosine and adenine methylation at **C**RGKG**A**TC motifs. A recently published overview of the epigenomic landscape of prokaryotes [67] did not mention any combinatorial cytosine and adenine methylations at common motifs. In contrast to G**A***T*C methylation, methylation of cytosine residues at **C**RGKG**A***T*C motifs was fractional and unequally distributed on the NC and FS chromosomes (Fig 6A-B). There are several candidate methyltransferases to perform this methylation. One orphan uncharacterized S-adenosylmethionine-dependent methyltransferase, BAA196NC_RS22825, is located on the pathogenicity plasmid and is significantly expressed in the culture FS. Further study is needed to check if this type of methylation is specific for the plasmid bearing *E. coli*. Three uncharacterized orphan methyltransferases, *yafS, yafE* and *yfiF*,are located on the chromosome and expressed in both cultures, NC and FS. Another methyltransferase,*yhdJ*, is believed to be responsible for type-II ATGC**A**T methylation [77]. This gene is expressed in both cultures; however, only one methylation at this motif was found in NC, and five instances of this methylation out of 1,706 available motifs were found in FS culture. Either this methyltransferase is inhibited post-transcriptionally that is common for prokaryotes [67], or this methyltransferase may have another recognition site.

Methylation of the recognition sites protects the host DNA from cleavage by cognate endonucleases; however, other authors reported that alternative patterns of methylation by orphan methylases may be naturally observed in long-term stationary phase and may prevent DNA replication [78, 79], participate in gene regulation [80] and even reduce resistance to antibiotics [81].

The global pattern of cytosine methylation at motifs **C**RGKG**A***T*C, C**C**AGGRAH and W**C**CCT*G*GYR shows more variations between the NC and FS chromosomes and plasmids. The role of CCWGG-methylation in gene regulation in both, eukaryotes and prokaryotes, has been reported in many publications [67]. In *E. coli*, CCWGG-methylation is a regulator of the stationary phase and stress response [69]. However, no phenotypic changes were reported for *E. coli* cells with under- and over-production of the Dcm methyltransferase suggesting that the number of methyltransferase molecules does not affect the methylation pattern [66]. The alternative distribution of methylated sites under the effect of FS-1 may contribute to a specific gene regulation fitting to survival with FS-1 but to the detriment of the resistance to antibiotics. The comparison of alterations in the global DNA methylation patterns of the NC and the FS-1 treated *E. coli* cultures with the differential gene expression did not allow deducing any statistically reliable links between presence/absence of methylated sites and up- or downregulation of the genes. Depending on the location of a methylated nucleotide within genetic regions recognized by transcriptional activators, inhibitors, non-coding RNA and/or DNA-dependent RNA polymerases, the effect of methylation may be ambivalent. Switching between alternative promoter regions and transcriptional start points may have no effect on the gene expression level but on properties and activities of the encoded proteins. Lower BM scores of cytosine methylation compared to adenine methylation may indicate that these sites were methylated only in a fraction of the bacterial population. It may complicate finding links between the alternative cytosine methylation and the gene regulation. On the other hand, it suggests a possible role of cytosine methylation in supporting necessary phase-variations and metabolic heterogeneity of bacterial populations.

Stable but alternative patterns of adenine and cytosine methylation in the NC and FS genomes discovered in this study demonstrates an importance of DNA methylation for adaptation of bacteria to the presence of iodine-containing nano-micelles in the medium and may be associated with the observed increase in susceptibility of the treated *E. coli* to gentamicin and ampicillin. However, it should be admitted that the altered pattern of epigenetic modifications cannot prove by itself that these variations play any specific role in the bacterial adaptation to halogen containing environments and/or in antibiotic resistance reversion. These hypotheses must be proved in additional studies focused on individual methylation sites and its role in gene regulation.

## Conclusion

Iodine is one of the oldest antibacterial medicines used by humans starting from its discovery by French chemist Bernard Courtois in 1811 [82]. Acquired resistance to iodine, in contrast to antibiotics, has never been reported. Application of iodine-containing nano-molecular complexes, such as FS-1, allows a broader use of iodine against antibiotic resistant pathogens. This study showed a profound alteration of the gene expression profile in the antibiotic resistant strain *E. coli* ATCC BAA-196 cultivated with a sub-bactericidal concentration of FS-1 that led to an increase in susceptibility to gentamicin and ampicillin. The analysis of differential gene regulation suggested that possible targets of iodine-containing particles are cell membrane fatty acids and proteins, particularly the cytochrome molecules, that leads to oxidative, osmotic and acidic stresses. Damaging of the cell wall and membrane structures increased penetrability of bacterial cells by toxic compounds and heavy metal ions accompanied with a reduction of the membrane potential due to the loss of intracellular potassium ions. Bacteria responded by an increased activity of anion pumps and the potassium uptake system, and by inhibiting of permeases to reduce the iodine flux. This general inhibition may complicate uptake of nutrients into cells, which already are stressed by damaged cell wall and cellular membrane structures. All these factors lead to an increased susceptibility to antibiotics that persists even after removal of FS-1 from the medium. Inability of a quick restoration of antibiotic resistance suggests an involvement of epigenetic and epistatic mechanisms in adaptation to FS-1. This hypothesis was supported by discovering a steadily altered DNA methylation pattern in the FS-1 treated *E. coli*. Another hypothesis that iodine may halogenate directly the bacterial DNA may be true for *S. aureus* but hardly for *E. coli*.

## Future Perspective

This work was performed as a part of a bigger project on studying the mechanisms of reversion of susceptibility to antibiotics by treatment of multidrug resistant bacteria with iodine-containing nano-micelles. This approach looks promising for combating antibiotic resistant infections. This phenomenon first was observed on multidrug resistant tuberculosis [11] but also was reproduced on *S. aureus* [9, 12] and *E. coli* (the current work). All these studies followed the same experimental protocol to allow comparison of the results. Further studies aimed at development of new drugs against antibiotic resistant nosocomial infections will be conducted on multiple clinical isolates, which were recently collected in clinics in Kazakhstan [83].

## Summary Points

- Treatment of the antibiotic resistant *E. coli* with iodine-containing nano-micelles increased its susceptibility to gentamicin and ampicillin.
- The profound alteration of the gene expression pattern of the FS-1 treated culture was observed.
- All aerobic metabolic pathways were strongly inhibited in the treated culture.
- Many nutrient uptake transporters were inhibited possibly as iodine entrance points.
- Specific gene regulation showed that the treated culture suffered from osmotic, oxidative and acidic stresses.
- Osmotic stress was caused by an increased penetrability of the bacterial cell wall damaged by iodine.
- Intensified anabolism of bacterial cells suffering from stresses and shortage of nutrients depleted their ability to withstand high concentrations of antibiotics.
- The treated culture showed the increased susceptibility to antibiotic after removal of the iodine-containing drug by cell washing;
- Altered nucleotide methylation profile implies a possible role of epigenetic mechanisms in gene regulation and antibiotic resistance reversion.

## Author Contributions

ISK – sequencing, data processing and visualization, manuscript writing and edition; ABJ – microbiological procedures, data processing and visualization, manuscript writing and edition; SVS – sequencing, data processing and visualization; TVK – microbiological procedures, data processing; ANM – microbiological procedures, HPLC, data processing and visualization; ZAI – microbiological procedures, HPLC, data processing and visualization; AII – project management and conceptualization, funding acquisition, manuscript writing and edition; MJ – data processing and visualization, manuscript writing and edition; ST – genome annotation; ONR – data processing and visualization, funding acquisition, manuscript writing and edition.

## Acknowledgements

We’d like to thank Dr. Guy Plunkett III, the Senior Scientist Emeritus at the University of Wisconsin–Madison, for his valuable consultations regarding the history of the strain *E. coli* ATCC BAA-196 used in this study as a model organism. Also, we’d like to thank the laboratories of Radiobiology and Radiochemistry (SCAID) for technical support in conducting radiobiological studies of DNA and the Control and analytical laboratory (SCAID) for technical support in conducting HPLC studies.

