## Supplementary material for "Transcriptomics and methylomics study on the effect of iodine-containing drug FS-1 on *Escherichia coli* ATCC BAA-196": Figure S2

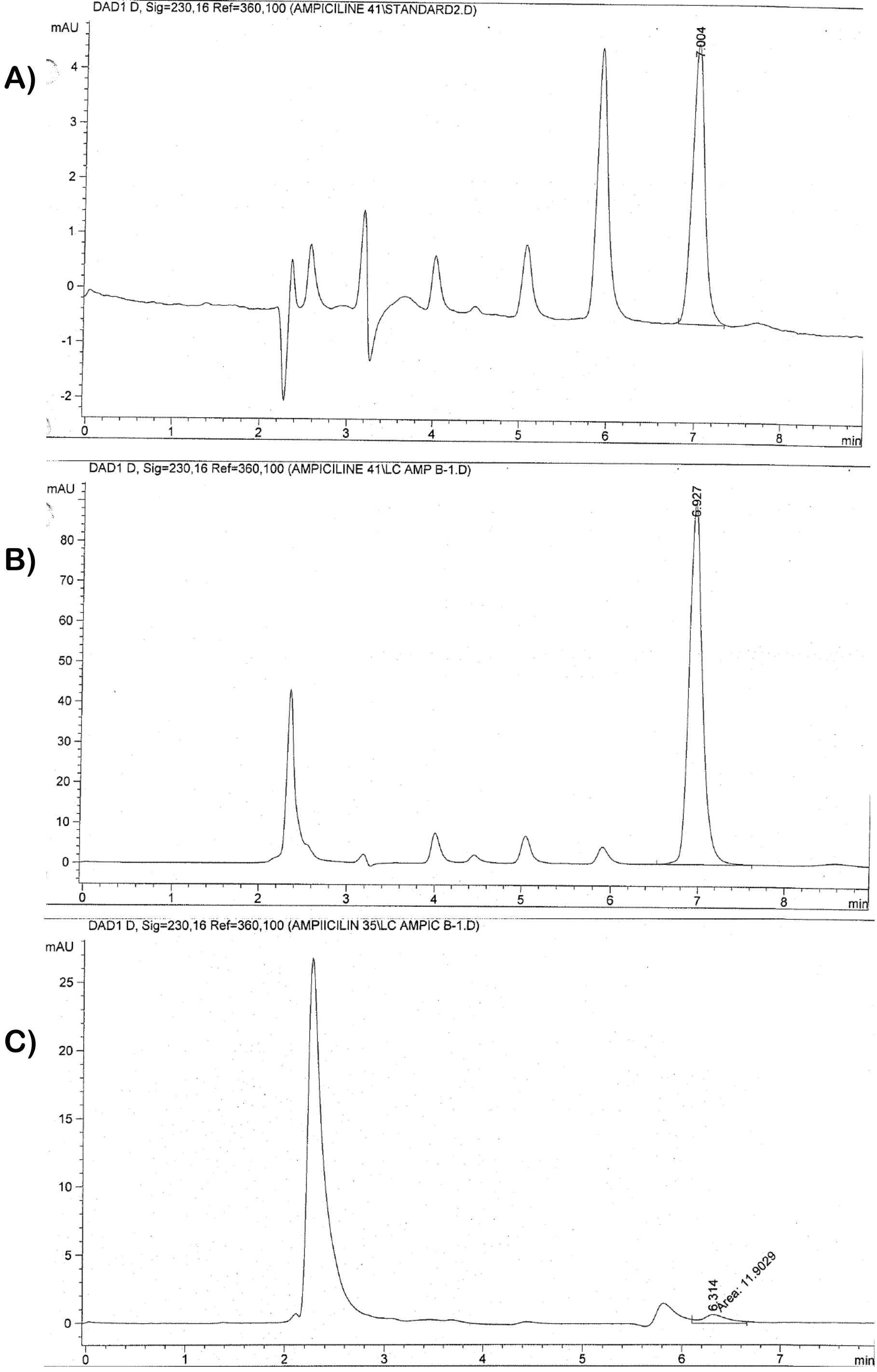


Supplementary Figures S1. HPLC peaks for ampicillin: A) standard antibiotic load 10 ppm (2.0 ng/ul); B) intracellular antibiotic in *E. coli* cells treated with the antibiotic and FS-1; C) intracellular antibiotic in *E. coli* cells treated with the antibiotic only. Axis X depicts elution minutes; axis Y - milli-Absorbance Units (mAU).
